# Discovery of dual thiobarbiturate-indole scaffold as a selective inhibitor targeting chikungunya virus nsP3 macrodomain through a cryptic binding pocket

**DOI:** 10.64898/2026.03.10.710793

**Authors:** Men Thi Hoai Duong, Tomi A.O. Parviainen, Aditya Thiruvaiyaru, Tero Ahola, Juha P. Heiskanen, Lari Lehtiö

## Abstract

The chikungunya virus (CHIKV) outbreak imposes a significant burden on healthcare systems and raises an urgent need for effective antiviral therapies. So far there are no specific drugs against CHIKV. A CHIKV macrodomain is critical for virulence and counteracts the host immune response, representing a promising antiviral drug target. Here, we describe small molecule inhibitors targeting the CHIKV macrodomain. Compound **1** (**MDOLL-0273**) was identified through a high-throughput screening using a fluorescence resonance energy transfer (FRET)-based assay, and its inhibitory activity was validated through multiple orthogonal assays. Compound **1** has a dual thiobarbiturate-indole scaffold and exhibits an IC_50_ of 8.9 µM. X-ray crystallography revealed that the inhibitor occupies an adenine binding site of the macrodomain and extends into a novel cryptic pocket. Notably, the inhibitor shows high selectivity for the CHIKV macrodomain over a panel of human and viral ADP-ribosyl binding and hydrolyzing proteins. Structure-activity relationship studies and medicinal chemistry efforts provide a promising starting point for further hit optimization.

## 1. Introduction

Chikungunya virus (CHIKV) belongs to the Alphavirus genus and Togaviridae family [1]. CHIKV is transmitted to humans via mosquitoes and has caused significant outbreaks worldwide [2–5]. The CHIKV infection is typically associated with acute symptoms such as severe fever and skin rash. In many cases, patients can develop chronic symptoms of severe arthralgia and neurological complications persisting for months and even years [6–9]. Recently, FDA has approved a live attenuated virus vaccine for CHIKV, and the vaccine will be primarily distributed in regions at high risk for severe outbreaks [10]. Effective antiviral treatments, however, are still under development.

CHIVK is an enveloped virus and has a nucleocapsid protecting a single-stranded, plus-sense RNA genome with 11.7 kilobases long [11,12]. A 5’ open reading frame (ORF) encodes a non-structural polyprotein (nsP1-nsP4) which is translated directly from the viral RNA genome (**Fig. 1A**) [11,13,14]. This non-structural polyprotein is cleaved into four individual proteins by a viral protease (nsP2) [11,15]. Four non-structural proteins further form a mature viral replication complex which is responsible for synthesizing full-length and subgenomic positive-strand RNAs [16,17].

**Figure 1.**
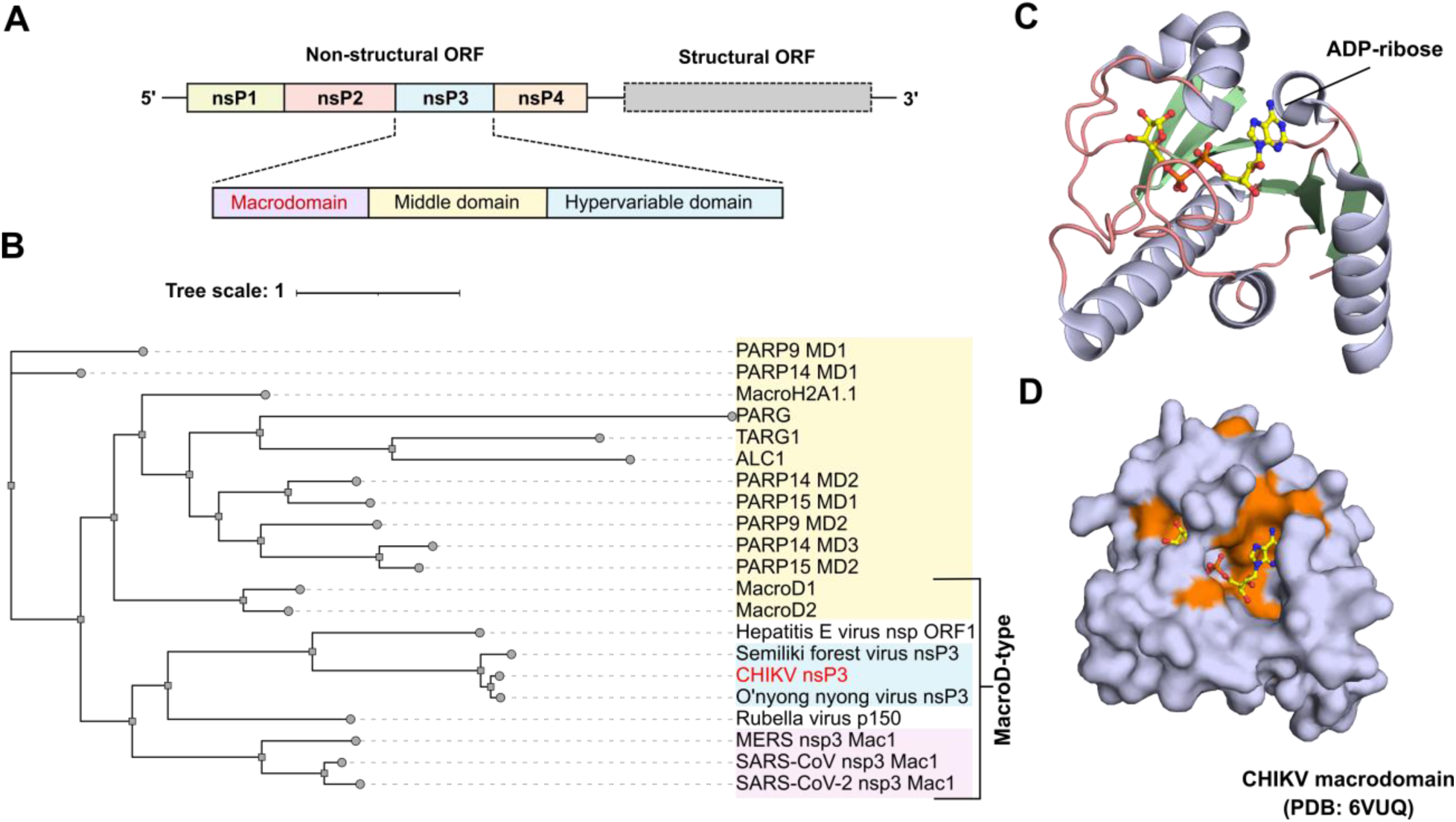
Conservation and structure of the macrodomain. **A)** Alphavirus genome architecture. Macrodomain is found in nsP3. **B)** Phylogenetic representation of macrodomains (MD) across representatives of alphaviruses (marked in blue), coronaviruses (marked in pink), and humans (marked in yellow). **C)** Structure of the CHIKV macrodomain in complex with ADP-ribose (PDB: 6VUQ) [40]. β-sheets are shown in green, α-helices are shown in light blue, loops are shown in dark salmon, ADP-ribose is shown in sticks and balls. **D)** Sequence conservation is mapped on the surface representation of the CHIKV macrodomain. Residues with similarity scores higher than 70% are shown in orange.

NsP3 has a macrodomain at the protein’s N-terminus (**Fig. 1A**). The macrodomain is an evolutionarily conserved protein fold found in all kingdoms of life from viruses to humans. Across viruses, macrodomains are encoded within the genomes of alphaviruses, coronaviruses, hepatitis E virus and rubella virus [18–20]. This domain is critical for viral replication and virulence and considered as a promising drug target for antiviral development [21–25]. During viral infection, several antiviral ADP-ribosyltransferases, for example, PARP9, PARP11, PARP12, PARP14, are induced by interferons [26–29]. In general, PARPs hydrolyze β-NAD^+^ and transfer either a single (mono-ADP-ribosylation, MARylation) or multiple (poly-ADP-ribosylation, PARylation) ADP-ribose units to proteins, nucleic acids, and small molecules. Most of the IFN-induced PARPs, however, are MARylating PARPs [18,30]. Viral macrodomain is characterized by its ability to remove these post-translational marks and counteract the host response [18,22,31]. The CHIKV macrodomain specifically removes the MAR moiety from aspartate and glutamate residues [32]. Mutations in the ADP-ribose binding site of the CHIKV macrodomain to abolish its MAR hydrolase activity prevented virus replication in mammalian BHK-21 cells or mosquito *Aedes albopictus* cells [33]. Moreover, a mutant with partially reduced MAR hydrolase activity replicated poorly in mammalian neuronal NSC-34 cell line and had reduced virulence in a neonatal mice model [33]. Furthermore, alphaviruses induce stress granule (SG) formation at the early stage of virus infection and the disassembly of SG was mainly caused by nsP3 [34]. Overexpression of the CHIKV macrodomain alone inhibited SG formation during CHIKV infection and the ADP-ribosylhydrolase activity of the macrodomain was also required to disrupt SG [35]. Together, targeting the CHIKV macrodomain represents a promising and underexplored antiviral therapeutic strategy.

Most macrodomains of the RNA viruses belong to a MacroD-type macrodomain subfamily, which includes two active human homologues: MacroD1 (also known as LRP16) and MacroD2 (**Fig. 1B**) [20]. In addition, the human genome encodes at least 14 other macrodomains that fall into distinct phylogenetic classes [36,37]. Structurally, the macrodomain has an α/β/α architecture, featuring a central β-sheet of six strands arranged in parallel and antiparallel, flanked by five surrounding α-helices and the ADP-ribose binds to a deep pocket formed by loops and turns (**Fig. 1C**) [19,38,39]. Due to a high degree of sequence conservation in the ADP-ribose binding site across viral and human macrodomains (**Fig. 1D, Fig. S1**), targeting viral macrodomains with small molecules for therapeutic intervention poses a risk of off-target effects, particularly on the human MacroD-type homologues, MacroD1 and MacroD2 [20]. Therefore, a critical assessment of inhibitor selectivity is essential in drug discovery efforts targeting viral macrodomains.

Despite the promising potential of the macrodomain as a drug target, only a few CHIKV macrodomain inhibitors have been identified, primarily through virtual screening and fragment-based screening coupled with X-ray crystallography. *In vitro* target-engagement validation and selectivity evaluation, however, are often lacking for hits identified through virtual screening [41–43]. In contrast, fragment-based screening with X-ray crystallography can rapidly provide binding modes for inhibitors. Zhang et al. employed this method and discovered a pyrimidone scaffold as a CHIKV macrodomain inhibitor [40]. This scaffold binds to the distal ribose binding site and has potential pan-antiviral activity. One compound from the series inhibited CHIKV replication in infected cells, with an EC_50_ of 23 μM, although its exact mechanism of action and possible off-target effect against human macrodomains remain unclear.

Here, we identified a novel, selective inhibitor targeting the CHIKV macrodomain using a developed fluorescence resonance energy transfer (FRET)-based high-throughput assay. The assay relies on the specific interaction between the CHIKV macrodomain and a MARylated peptide. Using this approach, we screened a SPECS chemical library of 30,335 compounds and identified compound **1** as a potential hit. Compound **1** has a dual thiobarbiturate-indole scaffold and exhibits an IC_50_ of 8.9 µM against the CHIKV macrodomain. Moreover, the compound inhibited the ADP-ribosyl hydrolase activity of the CHIKV macrodomain in a dot blot assay. We solved a complex crystal structure of the CHIKV macrodomain and the inhibitor to elucidate the binding mode. The inhibitor binds to the adenosine binding site and extends into a transient pocket formed by a flexible side chain of Arg1477. Notably, compound **1** demonstrated a high selectivity for the CHIKV macrodomain over a panel of human and viral ADP-ribosyl binding or erasing proteins. Furthermore, efforts in medicinal chemistry produced analogues with potency comparable to compound **1**. The synthetic route and structure–activity relationship analysis established in this study provide a valuable foundation for future hit optimization.

## 2. Results

### 2.1. Development and validation of the FRET-based high-throughput assay

FRET-based assays are widely used in biochemical studies to detect protein–protein, ligand-protein interactions and enable high-throughput screenings [23,44–52]. Here, we set out to establish the FRET-based high-throughput assay for the macrodomain of CHIKV nsP3. To generate a FRET pair, we recombinantly expressed mCitrine yellow fluorescence protein (YFP) fused with a C-terminal 10-mer peptide of Gi protein alpha subunit (GAP) and mCerulean cyan fluorescence protein (CFP) fused with the CHIKV macrodomain (CFP-CHIKV) [53,54]. The GAP-tag was then ADP-ribosylated at the C-terminus on a cysteine residue by the pertussis toxin subunit S1 (PtxS1) from the bacterium *Bordetella pertussis* to generate MARylated YFP-GAP [53–55]. In the FRET assay mixture, the interaction between the CHIKV macrodomain and the MARylated GAP brings the fluorescence donor CFP and the fluorescence acceptor YFP into close proximity and energy transfer from CFP to a nearby YFP occurs (**Fig. 2A**). As a result, the fluorescent emission of the CFP donor (at 410 nm) decreases while fluorescent emission of the YFP acceptor (at 527 nm) increases (**Fig. 2B**). We evaluate the binding interaction by calculating the ratiometric FRET (rFRET) values, which is defined as the ratio of the emission intensity at 527 nm to the emission intensity at 477 nm when the donor is excited at 410 nm. Since free ADP-ribose can bind to and inhibit the macrodomain activity, it is used as a negative control in the assay.

**Figure 2.**
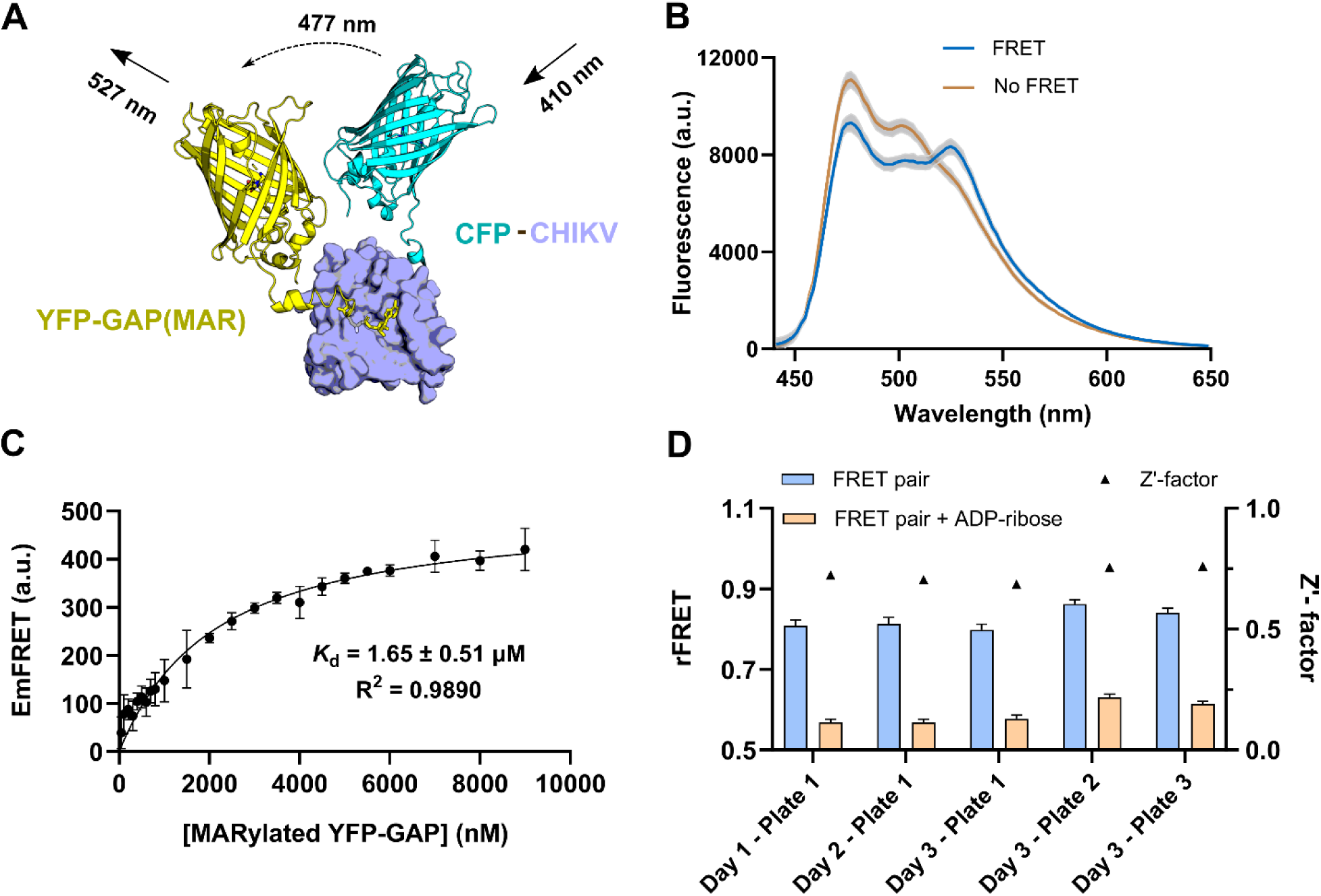
FRET-based high-throughput assay for the CHIKV macrodomain. **A)** Scheme illustrates the FRET-based assay of CFP-CHIKV and YFP-GAP(MAR). **B)** Fluorescence spectra of samples with FRET and samples without FRET (FRET pair in the presence of ADP-ribose). Data shown are the mean ± standard deviation of four technical replicates. **C)** Dissociation constant measurement of the FRET pair. Data shown are the mean ± standard deviation of four technical replicates. The *K*_d_ value is calculated from three independent experiments. **D)** Assay validation across five different 384-well format plates. rFRET data shown are the mean ± standard deviation of 176 replicates.

To measure the binding affinity *K*_d_ between CFP-CHIKV and YFP-GAP(MAR), we used the quantitative FRET-based technology, which is described by Song et al. [56] (**Fig. 2C**). The binding affinity was determined as 1.65 ± 0.51 µM, which is consistent with the dissociation constant between the CHIKV macrodomain and free ADP-ribose measured in the isothermal titration calorimetry (ITC) reported by Malet et al. (5.0 ± 0.4 µM) [57]. Considering the FRET-based *K*_d_ of 1.65 µM as a starting point, we optimized protein concentrations used in the high-throughput assay by varying the MARylated YFP-GAP concentration approximately around the *K*_d_ value to increase the assay sensitivity in the weak and moderate inhibitor identification (**Fig. S2**). Although conditions consuming lower protein concentrations are feasible, we selected 0.25 µM CFP-CHIKV and 1.25 µM MARylated YFP-GAP because both of the proteins can be purified and produced at high yield, and this setup likely minimizes potential compound interference with the raw fluorescence signals of the FRET pair compared to using lower protein concentrations.

To evaluate the quality of the developed assay for high-throughput screening, we performed the assay validation which assesses the reproducibility of FRET signals throughout multiple wells, multiple plates, and different days (**Fig. 2D**). A 384-well plate format was evaluated due to its intended application in the high-throughput screening campaign. In the plate, 176 replicates of the negative control (FRET pair in the presence of 200 μM ADP-ribose) and 176 replicates of the positive control (FRET pair only) were dispensed using an automated liquid handling system. Statistical parameters demonstrating the assay quality for high-throughput screening including signal-to-noise ratio (S/N), signal-to-background ratio (S/B), dynamic range and Z’-factor are summarized in **Table S1**. The Zʹ-factors evidenced for the separation between the negative and positive control were found to be between 0.68 and 0.76, which is higher than 0.5 and considered excellent for high-throughput screening [58].

### 2.2. High-throughput screening discovered compound 1 as a CHIKV macrodomain inhibitor

After validating the developed FRET-based assay quality, we employed the assay to screen a SPECS library of 30,335 small molecules against the CHIKV macrodomain. The screening was carried out in a singlet and at a single concentration of 30 µM compound. The average screening window coefficient Z’-factor across 94 screening plates was 0.84 ± 0.04 (**Fig. S3A**), demonstrating the reliable high-throughput screening. Because the assay itself is susceptible to fluorescence interference originating from compounds, we applied several filters to eliminate them. Firstly, we used two excitation wavelengths, at 410 nm and 430 nm, in measurements. In contrast to the fluorescent proteins, small molecules have sharper excitation and emission peaks, thereby the observed inhibitory activity caused by fluorescent compounds are expected to be different when the samples are excited at 410 nm and 430 nm wavelength. Secondly, if the fluorescence emission of the compound-treated sample was 20% higher or lower than that of controls, the compound was excluded from further analysis. After eliminating intrinsic fluorescence interfering compounds, we defined a hit limit to 19.5% which corresponds to three times the standard deviation from the mean value. The screening panel of 30,335 compounds is presented in **Fig. 3A**. The screening resulted in 122 hit compounds. After removing the common hits that were previously identified as primary hits but failed to pass through the validation step in the FRET-based high-throughput screening projects in our lab against other protein targets [23,49], we got 77 compounds as primary hits against the CHIKV macrodomain.

**Figure 3.**
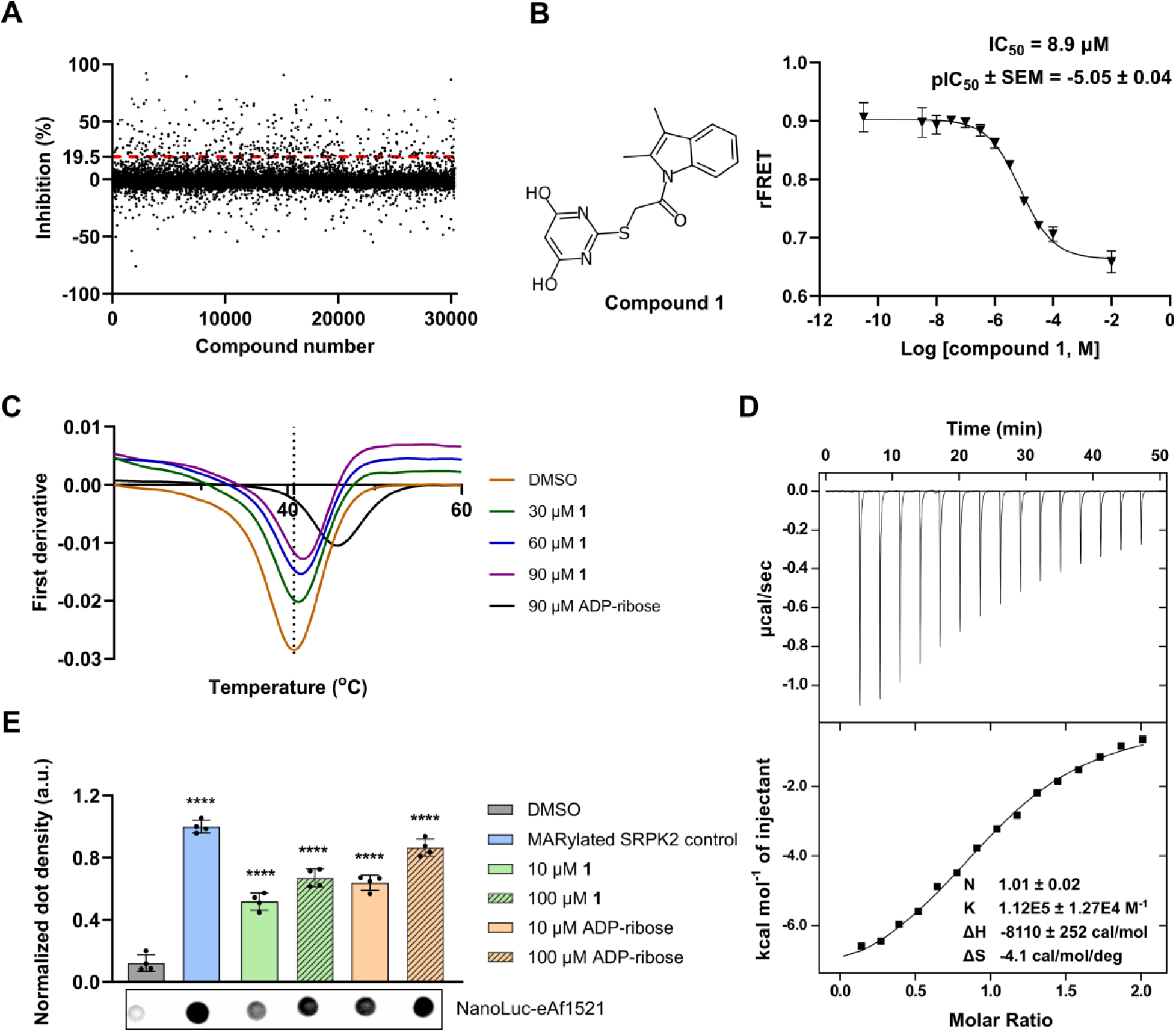
High-throughput screening and hit validation discovered compound **1** as a CHIKV macrodomain inhibitor. **A)** Screening panel of 30,335 compounds from the SPECS library against the CHIKV macrodomain. Data are shown as single measurements. The hit cut-off at 19.5% is shown as a red dash line. **B)** Dose-dependence inhibition curve of compound **1** against the CHIKV macrodomain using the FRET-based assay. An example curve is shown with mean ± standard deviation from four replicates. pIC_50_ is calculated from three independent experiments. **C)** Compound **1** caused the positive shift in the melting temperature of the CHIKV macrodomain in the dose-dependence manner. A representative curve of the three replicates is shown. The dot line represents the melting temperature of the DMSO control sample. **D)** ITC profile for the interaction between **1** and the CHIKV macrodomain. The upper panel presents raw injection heats of titrating **1** into the CHIKV macrodomain solution. The lower panel presents normalized integration data and a fitted curve. The curve was fitted using a single-site binding model. **E)** Dot blot assay showed the inhibitory activity of **1** against the hydrolysis activity of CHIKV macrodomain. Data shown are the mean ± standard deviation from four replicates, and representative dots are shown. ADP-ribose was used as a positive control. The statistical significance was calculated using one-way ANOVA with Dunnett’s multiple comparisons test of each condition against the DMSO control (**** p<0.0001).

To further exclude compounds that nonspecifically inhibited the FRET pair interaction, we employed a counter screen assay. We utilized a FRET-based assay developed for ankyrin repeat cluster 4 (ARC4) domain of tankyrase 2 (TNK2) and a tankyrase binding peptide motif [51]. Because the TNSK2 ARC4 peptide binding site is totally different from the ADP-ribose binding pocket of the macrodomain, it is unlikely that CHIKV macrodomain inhibitors can inhibit the TNSK2 ARC4 peptide binding activity. Additionally, we performed nano differential scanning fluorimetry (nanoDSF) in parallel with the counter screen. We identified 12 compounds that stabilized the CHIKV macrodomain in nanoDSF, 10 of which showed less than 20% inhibitory activity in the counter screen (**Fig. S3B-C**, **Table S2**). We purchased 7 of these compounds in powder form for further analysis as they were commercially available.

Dose-dependence inhibition analysis of 7 hits revealed that compound **1**, which bears a dual thiobarbiturate-indole scaffold, inhibited the ADP-ribosyl binding activity of the CHIKV macrodomain with an IC_50_ of 8.9 µM (pIC_50_ =-5.05 ± 0.04) (**Fig. 3B**). Compound **1** also stabilized the CHIKV macrodomain, demonstrating an increase in the protein melting temperature following the dose-dependence manner (**Fig. 3C**). We further confirmed the interaction between the inhibitor and protein using ITC. The result showed that the binding affinity was 8.93 ± 1.01 µM, the stoichiometry was 1.01, suggesting 1:1 ratio interaction between the inhibitor and the CHIKV macrodomain (**Fig. 3D**). To confirm the enzymatic inhibitory activity of compound **1**, we utilized the dot blot assay which measures the hydrolytic activity of the CHIKV macrodomain against glutamate/aspartate-linked MARylation. Consistent with the inhibitory activity observed in the binding assay, compound **1** also suppressed the hydrolase activity of the CHIKV macrodomain (**Fig. 3E**). The preliminary physicochemical parameters of compound **1** were evaluated to assess its drug-likeness using the SwissADME web tool [59] (**Table S3**). Key parameters including molecular weight, hydrogen bond acceptors, and hydrogen bond donors were calculated and found to fall within the acceptable range defined by Lipinski’s rule of five. Besides, the number of rotatable bonds and the topological polar surface area also satisfied Veber’s criteria, indicating a high likelihood of good absorption and membrane permeability. Calculated LogD at physiological pH (7.4) of 3.23 further indicates a good balance between the solubility and membrane permeability while still allowing room for structure-activity relationship optimization of the compound. Together, these results suggest that compound **1** represents a promising starting point in the CHIKV macrodomain inhibitor development.

### 2.3. Structure-activity relationship investigation

We next evaluated the inhibitory activity of compound **1**’s analogs to understand the structure-activity relationship supporting further hit optimization. Because there are no analogs of **1** found in the SPECS library, we first purchased 6 commercial analogs of **1**. The quality of analogs was then checked using high-performance liquid chromatography-mass spectrometry (HPLC-MS) to ensure that the compounds were intact. All 6 analogs have the modification at the thiobarbiturate ring, while the 2,3-dimethylindole ring is kept unchanged (**Table 1**). Removing two hydroxyl groups from the pyrimidine ring (compound **2**) completely abolished the compound’s inhibitory activity in the FRET-based assay. Similarly, substituting 4,6-dihydroxyl with either 4-methyl (compound **3**) or 4,6-dimethyl (compound **4**) resulted in compounds that exhibited markedly reduced activity. The results suggested that hydroxyl groups on the 4-and 6-positions of pyrimidine ring are fundamentally essential for the compound’s activity. Additionally, replacing the 4,6-dihydroxypyrimidine ring with smaller ring systems, including a 1,2,4-triazole ring (compound **5**) or *N*-methylimidazole ring (compound **6**), entirely eliminated the compound activity. Introducing a benzimidazole ring (compound **7**), however, resulted in an active compound with a decreased IC_50_ of 155 µM (pIC_50_ ± SEM =-3.66 ± 0.03), indicating that addition of a bulky hydrophobic moiety may partially compensate for the lack of hydroxyl groups.

**Table 1.** Commercial analogs with the variation at the thiobarbiturate ring.

| Compound ID | R | IC<sub>50</sub> (μM)<br>(pIC<sub>50</sub> ± SEM) |
| --- | --- | --- |
| 2 |  | NI |
| 3 |  | NI |
| 4 |  | 220 μM<br>(-3.66 ± 0.03) |
| 5 |  | NI |
| 6 |  | NI |
| 7 |  | 155 μM<br>(-3.81 ± 0.03) |

NI: No inhibition at 100 μM

Because the modifications at the thiobarbiturate ring decreased compound potency, we shifted our attention to the 2,3-dimethylindole ring and synthesized analogs altering this part. First, we established the synthesis route and resynthesized the original hit **1**. Our initial strategy was to first construct the (4,6-dihydroxypyrimidin-2-ylsulfanyl)acetic acid, and as the next step, to connect 2,3-dimethylindole through an amide linkage. In addition to enabling access to compound **1r**, this route would also allow for one-step variation of the indole moiety, facilitating the production of structural analogs. We managed to produce (4,6-dihydroxypyrimidin-2-ylsulfanyl)acetic acid with a reaction of 2-thiobarbituric acid and 2-chloroacetic acid but attempts to couple the produced compound with the indole nitrogen were unsuccessful. Therefore, we turned our attention to an alternative synthetic route, in which 2,3-dimethylindole was reacted with chloro acetylchloride attaching the chloroacyl-handle to the indole nitrogen in the first reaction step. Various attempts were made for this, but the lability of 2,3-dimethylindole in an acidic environment presented challenges due to the formation of hydrochloric acid side product. Luckily, the procedure reported by Mutschler and Winkler [60] was found to be viable. In this method, acyl chloride is added to a refluxing toluene solution containing an indole-derived starting material. Due to the low solubility of hydrochloric acid in hot toluene, the emerging hydrochloric acid is immediately expelled from the reaction solution and therefore does not interfere with the reagent or the reaction product. In the following reaction step, the thiolic proton of 2-thiobarbituric acid is first removed with aqueous sodium hydroxide, followed by addition of the intermediate chloroacylated 2,3-dimethylindole in tetrahydrofuran. This S_N_2 reaction is facilitated with the addition of tetrabutylammonium bromide (TBAB) as a phase transfer catalyst. With this synthetic strategy, we successfully resynthesized the high-throughput screening hit compound **1r** applying this strategy in the production of intermediates **8a**–**8d** and target molecules **9**–**11** (**Scheme 1**) as well. Compound **1r** had similar potency (IC_50_ = 12 µM, pIC_50_ ± SEM =-4.92 ± 0.02) compared to the commercial version compound **1** (IC_50_ = 8.9 µM). Starting with a small change in the indole ring, we substituted 2,3-dimethylindole with 3-methylindole (compound **9**). To our surprise, compound **9** was completely inactive. It is quite rare but possible that a methyl group can have a profound effect on the activity of a compound [61]. Substituting 2,3-dimethylindole with 1,2,3,4-tetrahydrocarbazole (compound **10**) resulted in 4 times lower potency (IC_50_ = 39 µM, pIC_50_ ± SEM =-4.41 ± 0.04). Notably, introducing an additional phenylene fused to positions 4 and 5 of the 2,3-dimethylindole ring (compound **11**) led to similar potency compared to the original hit (IC_50_ = 10 µM, pIC_50_ ± SEM =-4.98 ± 0.04). We anticipated that the indole binding site is spacious and can accommodate a planar structure without steric clashes.

**Scheme 1.**
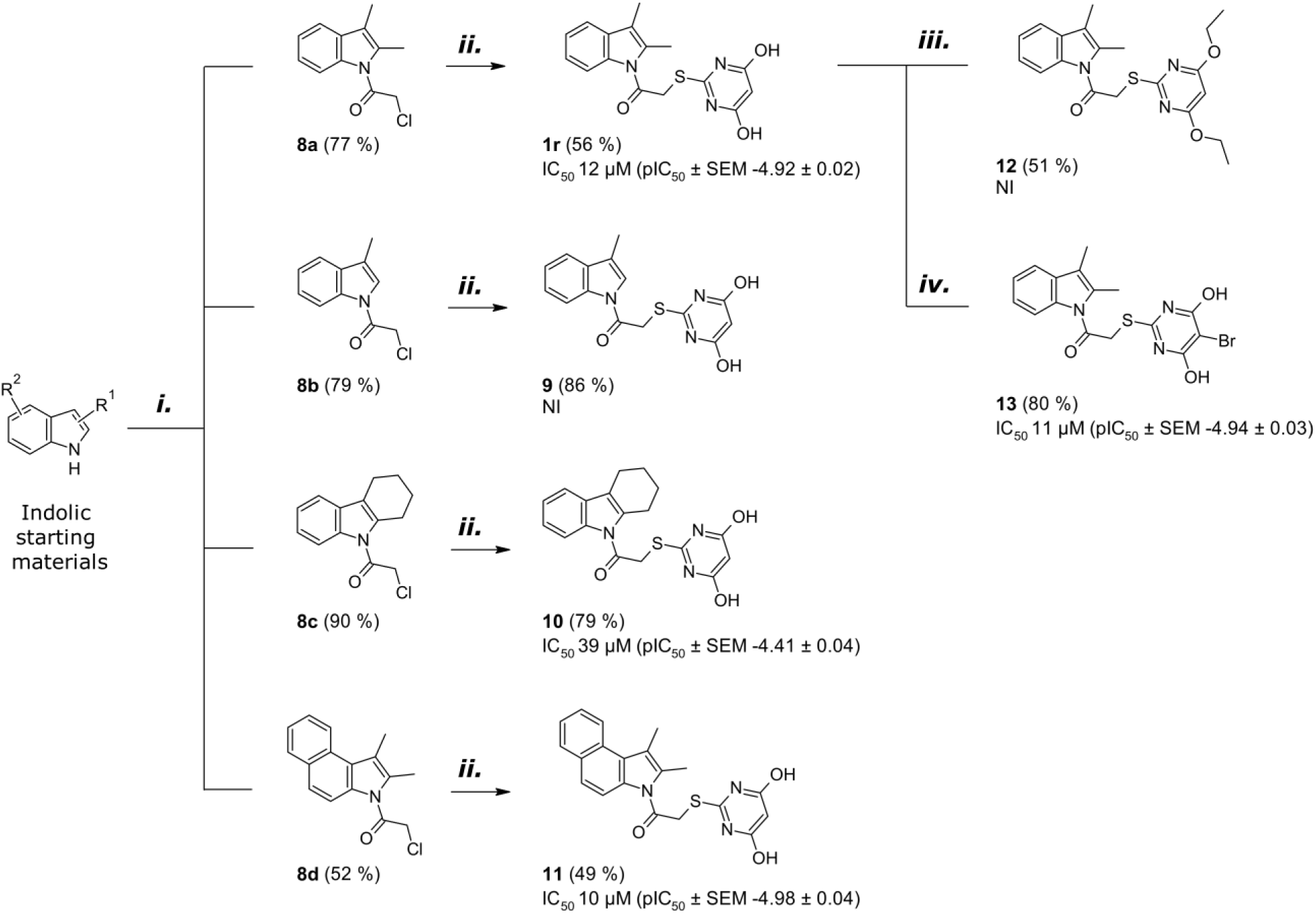
Reagents and conditions: (i) corresponding indole, toluene, reflux, then chloroacyl chloride, overnight (16–22 h); (ii) NaOH, H_2_O, 2-thiobarbituric acid, compounds **8a**–**8d**, THF, 50–55 °C, ∼3 days; (iii) diethyl sulphate, K_2_CO_3_, Ar, 80 °C, 4 h; (iv) NBS, DMSO, rt, 2½ h.**2.4.**

With synthesized **1r** at hand, we created two compounds extending the structure-activity relationship analysis of the commercial pyrimidine analogs (**Table 1**). We generated a diethylated analog of 4,6-dihydroxypyrimidine ring (compound **12**), treating compound **1r** with diethyl sulphate. Similar to the commercial analogs, **12** was completely inactive. Introducing a bromine atom to position 5 of the thiobarbiturate ring resulted in an analog (compound **13**) with potency comparable to the original hit (IC_50_ = 11 µM, pIC_50_ ± SEM =-4.94 ± 0.03) and similar cLogD value at pH 7.4 of 3.77. This further confirmed the requirement of the hydroxyl groups for an efficient binding while indicating that some additional modifications to the thiobarbiturate ring may be considered in the future.

### Compound 1 occupies the adenosine binding site and extends to a transient pocket of the CHIKV macrodomain

To understand how **1** interacts with the CHIKV macrodomain, we solved a crystal structure of the complex. **1** was soaked into apo CHIKV macrodomain crystals. The protein was crystallized in space group *P3_1_* with four copies in an asymmetric unit, and the structure model was refined to 1.65 Å (**Table S4**). The CHIKV macrodomain sequence was numbered according to a Uniprot entry Q8JUX6 (A1334-T1493) and this numbering is used in the below analysis. Consistent with the deposited structures on protein data bank, the CHIKV macrodomain structure comprises six stranded-β sheets sandwiched by two α-helices and three α-helices. According to an available deposited structure (PDB: 3GPO) [57], ADP-ribose is stabilized by an extensive direct and water-mediated hydrogen bonding network as shown in **Fig. 4A**. **1** was modelled in the positive electron density found in all four protein chains and occupied the adenosine binding site (**Fig. 4B**), which is consistent with its ADP-ribose competitive binding activity shown in the FRET-based assay. Notably, while the thiobarbiturate ring overlapped with the adenosine site, the 2,3-dimethylindole ring was perpendicularly extended further to a transient pocket formed by Arg1477 and Asp1343 (**Fig. 4C-E**). In contrast to the ADP-ribose complex structure where Arg1477 was flexible and adopted multiple conformations, the residue was stably positioned in the complex with compound **1** (**Fig. S4**). The indole ring stayed between Arg1477 and Asp1343 in a sandwich-like configuration and was stabilized by the cation-π interaction with Arg1477 and the anion-π interaction with Asp1343. Additionally, 2-methyl substituent pointed toward a hydrophobic site formed by the side chains of Ala1369, Val1366, and Ile1344 further stabilizing the complex (**Fig. 4F**). This unique binding mode of the 2,3-dimethylindole ring can explain the reduced potency observed in analogue **10**. The 1,2,3,4-tetrahydrocarbazole ring of compound **10** may possibly clash with the nearby Ala1369 and disturb the local electrostatic environment around Arg1477 and Asp1343, leading to unfavorable binding and weaker potency.

**Figure 4.**
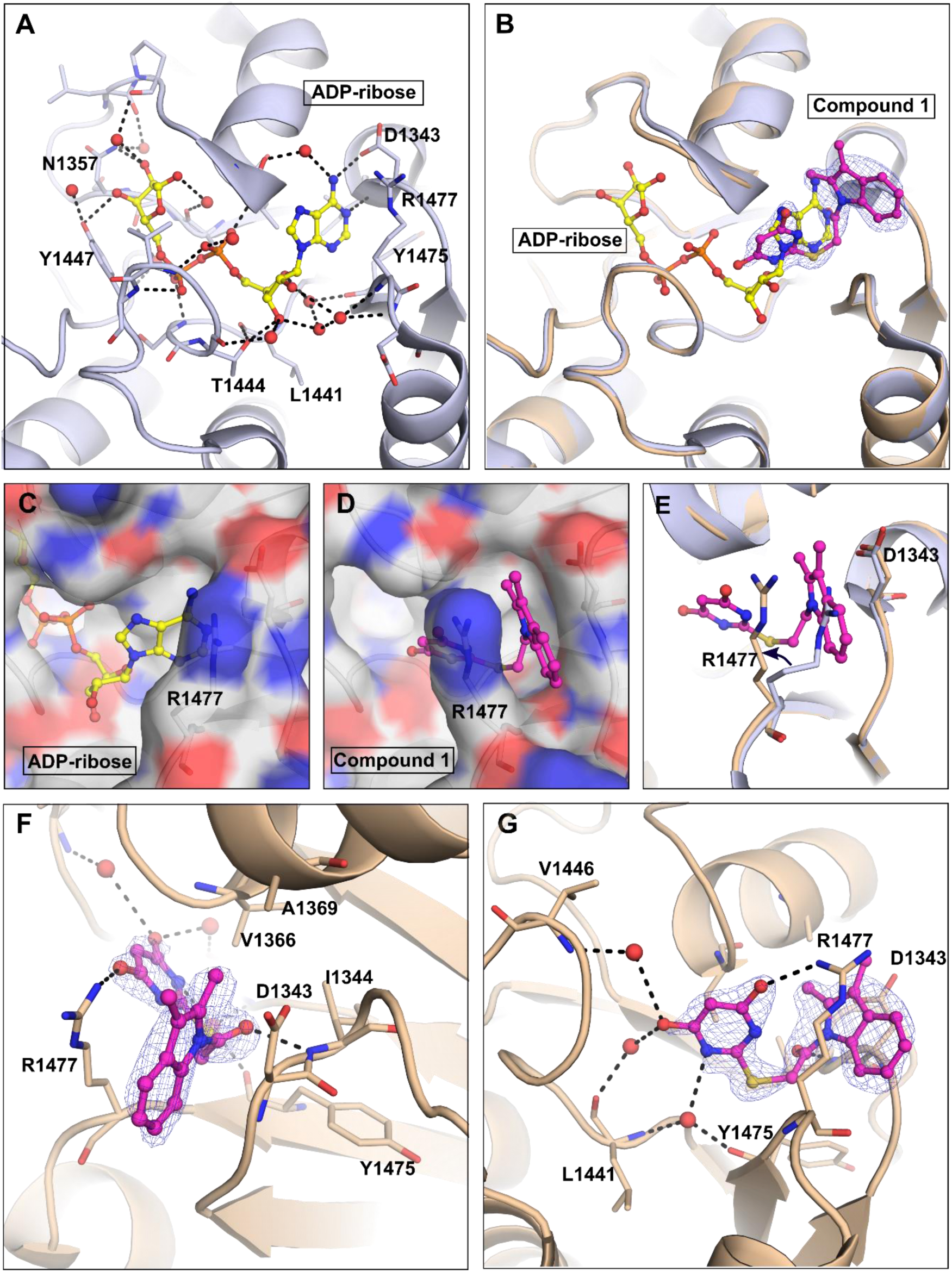
Binding mode of **1** in the ADP-ribose binding pocket of the CHIKV macrodomain. **A)** ADP-ribose binds to the CHIKV macrodomain and is stabilized through a network of hydrogen bonding interactions (PDB: 3GPO) [57]. The protein is represented as a light blue cartoon. ADP-ribose is displayed as a yellow stick-and-ball model. Water molecules are displayed as red spheres. Hydrogen bonds are shown as black dash lines. **B)** Overlay the CHIKV macrodomain structure in complex with ADP-ribose (PDB: 3GPO) [57] and with compound **1** (sand cartoon, PDB: 9TDQ). Compound **1** is displayed as a magenta stick. The electron density map of **1** is shown as a blue mesh contoured to 1σ. **C-D)** Surface presentation of the CHIKV macrodomain in complex with ADP-ribose **(C)** and compound **1 (D)** highlights a transient pocket between Arg1477 and Asp1343. **E)** Arg1477 is flexible and moves away to accommodate **1**. Arg1477 is displayed as a light blue stick (in ADP-ribose complex structure) and a sand stick (in compound **1** complex structure). **F-G)** Two different views show binding interactions of **1** and the CHIKV macrodomain. The residues involved in the binding are displayed as sticks. Water molecules are displayed as red spheres. Hydrogen bonds are shown as black dash lines. The first view (**F**) focuses on the interaction between the indole ring and the transient pocket. The second view (**G**) focuses on the interaction between the thiobarbiturate ring and the adenosine binding site.

In the adenosine binding site, the hydroxyl groups of the 4,6-dihydroxypyrimidine ring formed a hydrogen bond with the side-chain amide of Arg1477, as well as water-mediated hydrogen bonds with the backbone of Val1446 in the phosphate binding loop and Leu1441 in the proximal ribose binding site (**Fig. 4G**). The loss of these hydrogen bonds explains the loss in the inhibitory activity of commercial analogs in which the hydroxyl groups in positions 4 and 6 were either removed or substituted with methyl groups (**Table 1**). Additionally, the amide group in the pyrimidine ring formed water-mediated hydrogen bonds with the backbone of Leu1441 and Tyr1475 in the proximal ribose binding site. Moreover, the carbonyl oxygen atom of the 2-sulphidoacyl linker formed a hydrogen bond with the backbone amide of Ile1344 in the adenine binding site (**Fig. 4F**). Notably, the residues Val1446, Leu1441, Tyr1475, and Ile1344 were also involved in hydrogen-bonding interactions with ADP-ribose (**Fig. 4A**). Introducing a bromine atom at position 5 of the thiobarbiturate (compound **13**) could potentially form a dipole-dipole interaction with a nearby water molecule. However, this interaction appears to have minimal impact on the binding affinity as the compound’s potency was not improved.

### 2.5. Compound 1 is selective for the CHIKV macrodomain over other macrodomains

Compounds targeting the viral macrodomain for infection treatment naturally carry a high risk of the off-target effect since humans bear 16 macrodomains that share the conserved fold, of which 13 macrodomains are active. To assess this potential risk, we next evaluated the selectivity of **1** by profiling it against a panel of all active human macrodomains. In addition to macrodomains, we also accessed the possibility of the off-target effect against an ADP-ribosyl hydrolase ARH3, which has an unrelated structure fold and is therefore less likely to interact with the compound, although it is capable of removing the ADP-ribose mark on serine. Besides, compound’s potential inhibitory activity against coronavirus macrodomains and another alphavirus macrodomain was also evaluated. Overall, the result showed that compound **1** demonstrated high selectivity for the CHIKV macrodomain over the tested panel (**Fig. 5A**). It showed only weak inhibition for SFV, PARG, ALC1, and PARP14 macrodomain 1, with approximately 40% inhibition observed at 100 µM. The cross-activity against SFV is expected as SFV and CHIKV both belong to the alphavirus genus and share relatively higher sequence identity (76.74%) compared to other macrodomains. Although very weak inhibitory activity was observed against PARG, ALC1 and PARP14 macrodomain 1, possible cross-activity against those proteins should be considered when designing the new analogues in the future.

**Figure 5.**
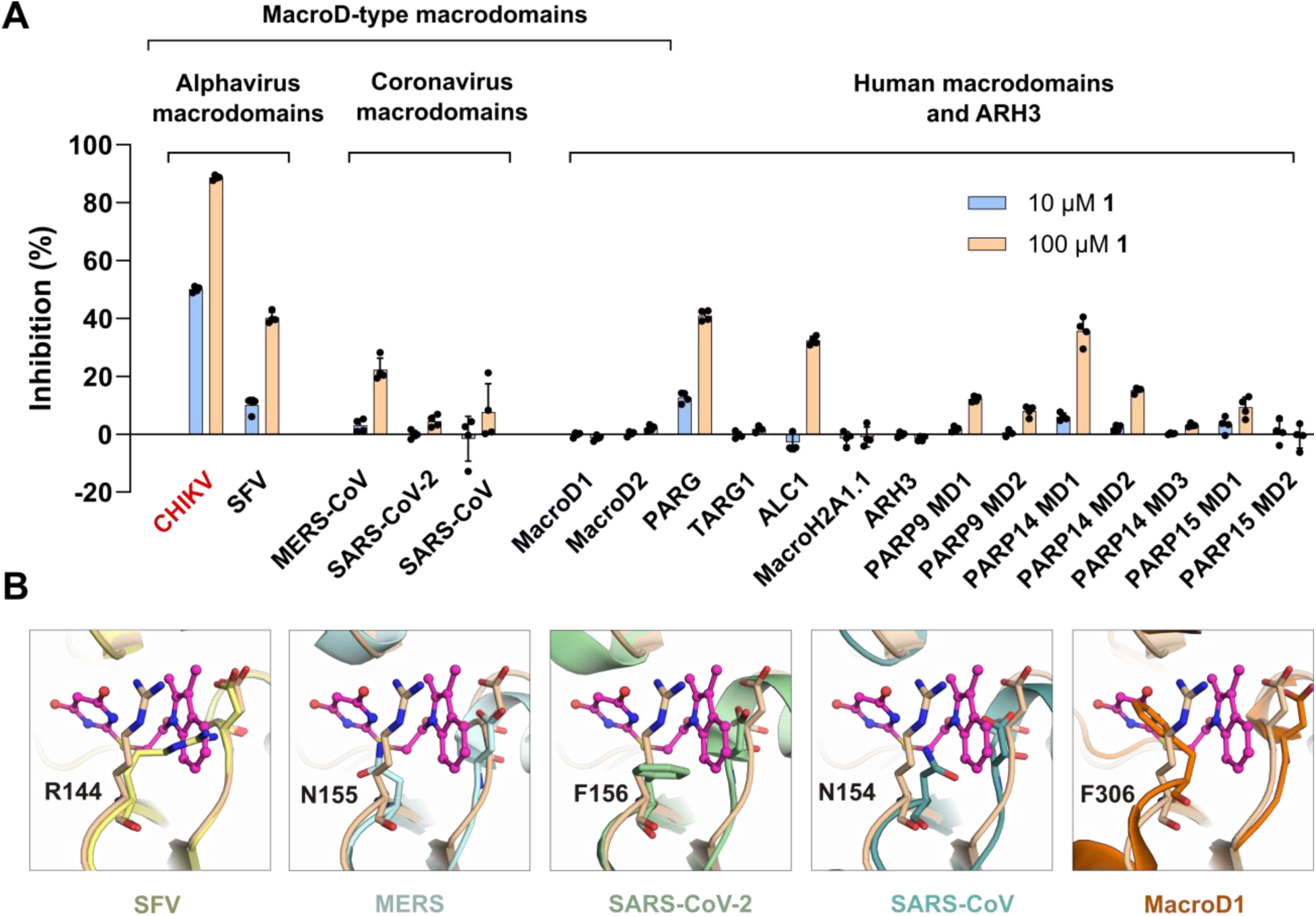
Profiling panel of **1** against viral macrodomains and human macrodomains (MD). The inhibition activity was measured using FRET-based assays as described earlier [53]. Data are presented as the mean ± standard deviation from four replicates. The individual data points are shown. The crystal structures of the MERS macrodomain (PDB: 5HOL) [62], SARS-CoV-2 Mac1 (PDB: 8TV7) [23], human MacroD1 (PDB: 2X47) [63] and alphafold predicted models of SFV macrodomain and SARS-CoV Mac1 are superimposed to the structure of the CHIKV macrodomain:**1** complex (PDB: 9TDQ). The CHIKV macrodomain is presented as sand cartoon and compound **1** is presented in the magenta stick-and-ball model.

Multiple sequence alignment and structural comparison revealed a key difference at the residue Arg1477. While this positively charged arginine is found unique for the CHIKV macrodomain and is shared with at least two other alphaviruses SFV and O’nyong nyong virus, the hydrophobic and aromatic residues are commonly found at the corresponding position in other examined macrodomains (**Fig. S1**). A closer look to the MacroD-type subclass revealed that the corresponding residue is phenylalanine in SARS-CoV-2 and MacroD1, whereas it is asparagine in MERS and SARS-CoV (**Fig. 5B**). While Arg1477 is flexible and allows forming a transient pocket to accommodate the inhibitor, phenylalanine is rigid and may introduce steric clashes with the compound. Replacing arginine with asparagine could change the local electrostatic environment at this position, which potentially makes it unfavorable for compound interaction. To confirm the contribution of Arg1477 to the activity and selectivity of the inhibitor, we mutated this residue to either phenylalanine or alanine. Notably, the compound activity exhibited a significant loss against both CHIKV macrodomain R1477F and R1477A mutants with the potency dropping by one order of magnitude (**Fig. S5**). Arg1477 is therefore evident as a key determent responsible for the inhibitor activity and selectivity, and exploiting the cryptic binding site formed by this arginine offers a strategy for developing selective inhibitors targeting alphavirus macrodomains while avoiding undesired cross-reactivity with human macrodomains.

### 2.6. Cell-based assay

Next, we studied the effect of **1** on CHIKV during virus infection under cell culture conditions. We used an attenuated strain of CHIKV, 181/25 [64], which was further modified to express a NanoLuc reporter gene [65]. The macrodomain sequence of this strain is 100% identical to that of the recombinant CHIKV macrodomain used in our biochemical assays. The cells were incubated with compound **1** at three concentrations, 12.5 µM, 25 µM, and 50 µM, at which we evaluated CHIKV infection as well as cell viability. There was no significant (p>0.05) cytotoxic effect of **1** at these concentrations based on the ATP level measurements in both BHK-21 and HEK-293T cells (**Fig. S6**). However, under cell culture conditions we could also not observe significant (p>0.05) reduction in CHIKV-nLuc replication in either BHK-21 or HEK-293T (**Fig. 6**). Considering that the inhibitor’s potency (IC_50_ = 8.9 μM) may be insufficient to allow us to observe any measurable effects in cell-based assays, we would need to further optimize this hit compound before proceeding additional cellular studies.

**Figure 6.**
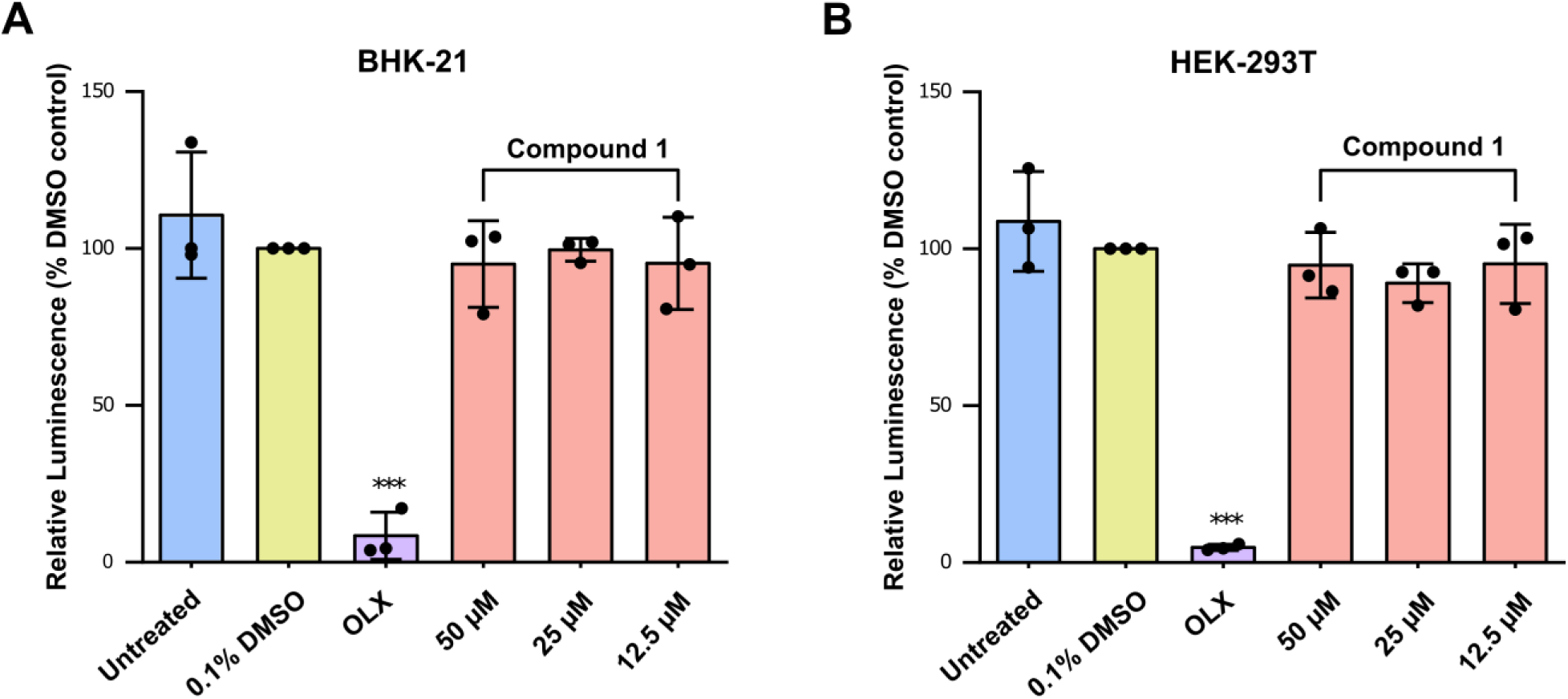
Effect of compound **1** during CHIKV infection. The cell lines **A**) BHK-21 or **B**) HEK-293T were infected with CHIKV-nLuc in the presence of **1** at the indicated concentrations at MOI 0.01 or MOI 0.05 respectively. The response of the inhibitor on the CHIKV-nLuc replication was determined by measuring the NanoLuc luciferase activity at **A**) 16 h or **B**) 20 h post-infection. The reporter activity was normalized with respect to the DMSO control and represented as a mean of three independent biological repeats (each carried out in quadruplicate) ± standard deviation. The statistical significance was calculated using one-way ANOVA with Dunnett’s multiple comparisons test of each condition against the DMSO control; *p<0.05; **p<0.01; ***p<0.001. 0.5 µM obatoclax (OLX) was used as a positive control.

## 3. Discussion

Here, we developed a robust, high-throughput FRET-based binding assay for identifying CHIKV macrodomain inhibitors. The assay is simple, automated, cost-efficient, and does not require specialized reagents. It was scaled to screen a large compound library and can be further expanded for even larger collections. Currently, only a few efforts to develop CHIKV macrodomain inhibitors have been reported, all of them rely on virtual screening or fragment-based X-ray crystallography [40–43,66]. Although computational approaches offer high-throughput and cost-efficient strategies for identifying potential small-molecule inhibitors, they often lack direct experimental evidence confirming binding to the CHIKV macrodomain. Fragment screening by X-ray crystallography can rapidly reveal structural insights into small-molecule binding modes, but it is typically limited by the low binding affinities of fragment hits and the low-throughput format. Our biochemical assay complements these existing methods and provides a valuable tool to accelerate high-throughput screening for novel inhibitors.

Using this assay, we identified **1** as a selective CHIKV macrodomain inhibitor. The inhibitor has an IC_50_ of 8.9 µM in the FRET-based assay. Moreover, the inhibitor significantly suppressed the enzymatic activity of the CHIKV macrodomain against glutamate/aspartate-linked MARylated protein. This inhibitor has a dual thiobarbiturate-indole scaffold. Barbituric acid derivatives have a long history of usage as sedatives and anesthetics, but these typically lack the sulfur atom in position 2 (most notable exception being thiopental) of the barbiturate ring, bearing an oxygen atom instead. In addition, their substituents are typically placed in carbon 5 or to the nitrogen atom(s). Substituted indole, on the other hand, is a common motif in many pharmaceuticals, e.g. in antiviral and pain management medications [67].

To date, only two studies have reported CHIKV macrodomain inhibitors with clear binding modes, both utilizing fragment screening approaches [40,66]. The identified fragments either occupy the ADP-ribose binding site or stay at the novel transient pockets formed by the flexible Arg1477 residue. Notably, compound **1** spans both the cryptic pocket and the adenine-binding region, suggesting a distinct binding mode. Moreover, although the residues within the ADP-ribose binding site are highly conserved, the Arg1477 is found unique for the alphavirus macrodomains and can be exploited to gain selective inhibition for the CHIKV macrodomain over human counterparts.

The lack of inhibitory effects of compound **1** in cell culture is likely due to its moderate potency. Although a cellular thermal shift assay (CETSA) could be used to assess target engagement in cells, the modest thermal shift observed in nanoDSF indicates that CETSA is unlikely to be informative at this stage. Enhancing compound potency is therefore essential before going with antiviral studies. Given that a pan-antiviral 2-pyrimidone-4-carboxylic acid fragment was identified to occupy the distal ribose site of the CHIKV macrodomain by another research group [40], incorporating this fragment into the dual thiobarbiturate–indole scaffold described in this work may offer a promising approach to enhance inhibitor potency while preserving its selectivity. Suitable linkers could include aromatics, such as benzylic structures or their bioisosteres. Extension of the inhibitor structure could be achieved, for example, through coupling chemistry, as the 5-position of the thiobarbiturate moiety is readily brominated. In addition, the inherent nucleophilicity at the 5-position of the thiobarbiturate could also be utilized for this purpose. The 2-pyrimidone-4-carboxylic acid fragment could then be attached to the linker via an amide bond formed through its carboxylic acid functionality, using an amine-containing linker and various methods available for amide bond formation. However, appropriate linkers must be carefully designed and optimized to have a certain degree of flexibility and preserve the binding geometries of both pharmacophores. Nevertheless, the structure-activity relationship analysis and established chemical synthesis route presented in this work offer valuable guidance for further hit optimization effort.

## 4. Materials and Methods

### 4.1. Bioinformatics

Protein sequences of macrodomains were retrieved from the UniProt database. Multiple sequence alignment was performed using the PROMALS3D online server [68] and visualized with ESPript 3 [69]. The similarity scores were calculated using the % Equivalent scheme in ESPript 3. The phylogenetic tree was constructed using the IQTREE server with the LG substitution model and maximum-likelihood method [70,71]. The tree with the highest likelihood is visualized using the iTOL web server [72].

### 4.2. Cloning, protein expression and purification

The plasmid of the CHIKV macrodomain (Uniprot Q8JUX6, from residue A1334 to T1493) fused to pNIC-CFP vector (Addgene: 173083) is available in our lab from previous work [53].

YFP-GAP plasmid is available in our lab from previous work and deposited in Addgene: 173080 [53].

MARylated YFP-GAP was produced as described earlier [53,54].

The CFP-CHIKV plasmid was transformed to *E.coli* BL21 (DE3) competent cells and cells were grown overnight at 37 °C on an agar plate containing 50 µg/ml kanamycin. Next day, several colonies were inoculated in Luria-Bertani broth medium and cultured overnight at 37 °C in a shaking incubator. The seed culture was then added to the main culture following 1:100 ratio. One liter of Terrific Broth autoinduction media including trace elements containing 8 g/L glycerol and 50 μg/mL kanamycin was used as the main culture. Cells were cultured at 37 °C in the shaking incubator for approximately 2-3 h until the optical density at 600nm (OD_600_) reached 1.8. Subsequently, cells were further cultured at 18 °C for approximately 20 h to allow protein production. Cells were harvested and the pellet was resuspended in lysis buffer (50 mM HEPES, pH 7.5, 500 mM NaCl, 10 mM imidazole, 10% (v/v) glycerol, and 0.5 mM TCEP) containing 0.1 mg/mL lysozyme, 0.02 mg/mL DNase, 0.1 mM protease inhibitor Pefabloc (Sigma-Aldrich, USA).

Cells were lysed using sonication, and the lysate was then centrifuged at 16,000 rpm for 45 min at 4 °C. The lysate was filtered through 0.45 µm membrane filter and then loaded onto a 5 mL immobilized metal affinity chromatography (IMAC) column charged with Ni2^+^ at the flow rate of 2 ml/min. The column was then washed extensively with lysis buffer followed by wash buffer containing 25 mM imidazole. Protein was eluted with an elution buffer containing 150 mM imidazole. Eluted protein was concentrated using 30 kDa molecular weight cut off (MWCO) concentrator and loaded onto a Hiload 16/600 Superdex 75 (GE Healthcare, Sweden) gel filtration column. The running buffer contains 30 mM HEPES, pH 7.5, 300 mM NaCl, 10% (v/v) glycerol, and 0.5 mM TCEP. The purified protein was concentrated, divided into 50 μL aliquots and flash frozen in liquid nitrogen to be stored at-70 °C. For non-tagged CHIKV-nsP3 macrodomain used in biophysical assays and crystallization, an additional step of TEV protease treatment was performed, followed by reverse IMAC to remove 6His-tagged CFP and any impurities.

Other proteins used in the profiling were expressed and produced as previously described [53].

### 4.3. Dissociation constant (*K*_d_) measurement

The binding affinity of the CHIKV macrodomain and MARylated peptide was measured using quantitative FRET technology [56]. CFP-CHIKV concentration was kept constant at 500 nM. MARylated YFP-GAP concentration varied from 50 nM to 9000 nM. Echo 650 acoustic liquid dispenser (Beckman Coulter Life Sciences, Indianapolis, IN, USA) was used to dispense proteins to a black and flat-bottom ProxiPlate-384 F microplate (Revvity, Waltham, MA, USA). Mantis liquid handler (Formulatrix, Bedford, MA, USA) backfilled the reaction wells with the buffer containing 10 mM BTP, pH 7.0, 3% (w/v) PEG 20,000, 0.01% (v/v) Triton X-100, and 0.5 mM TCEP to 10 μL per well. The fluorescence was measured using Spark plate reader (Tecan, Männedorf, Switzerland). The samples were excited by 430 nm light and the fluorescence emission at 477 nm and 527 nm were measured. Additionally, the emission at 527 nm was measured upon excitation at 477 nm. The analysis of the data was done in GraphPad Prism 10 as described by Song et al. [56].

### 4.4. FRET assay validation

A 384-well plate was set up with 176 wells designated for positive controls, 176 for negative controls, and 32 blanks. The blank wells contained only the buffer. For the positive control, each sample well contained the FRET pair of 0.25 µM CFP-CHIKV and 1.25 µM MARylated YFP-GAP. In contrast, the negative controls had the same FRET components with the addition of 200 µM ADP-ribose. The well volume is 10 µL. To assess consistency, five plates were used: one tested on day one, another on day two, and the remaining three plates tested on day three.

### 4.5. High-throughput screening

A library of 30,335 compounds from SPECS consortium collection was screened against the CHIKV macrodomain. The 384-well plates containing 30 nL of 10 mM compound stock per each well were supplied by the Institute for Molecular Medicine, Finland (FIMM). The screening was carried out in a singlet and at 30 μM compound concentration. Each sample well contained 10 μL of the FRET mixture consisting of 0.25 μM CFP-CHIKV and 1.25 μM MARylated YFP-GAP.

### 4.6. Counter screen

We used the FRET-based assay developed for the ARC4 domain of TNKS2 and TBM for counter screen [51]. The hit compounds were tested in duplicates at a concentration of 30 μM. The pre-plated compounds (30 nL of 10 mM stock) were purchased from FIMM. 10 μL of the FRET mixture consisting of 100 nM CFP-TNKS2 ARC4 and 200 nM YFP-TBM was added to the plates. 1 M GdnHCl was used as a negative control to disrupt the protein-peptide interaction.

### 4.7. Nano differential scanning fluorimetry

0.3 mg/ml of the CHIKV macrodomain was mixed with hits in the buffer containing 10 mM HEPES, pH 7.0, 100 mM NaCl, and 0.5 mM TCEP. The compound was tested at different concentrations of 30 µM, 60 µM, and 90 µM. Final DMSO concentration was 2% (v/v). 90 µM ADP-ribose was used as a positive control. Measurements were performed using Prometheus NT.48 (nanoTemper, Germany). The melting curves were recorded with a 1°C increment per min.

### 4.8. Potency measurements

Serial half-log dilutions of hit compounds from 100 µM to 0.003 µM were tested. 10 µL mixture of 0.25 μM CFP-CHIKV and 1.25 µM MARylated YFP-GAP (final concentration) in the FRET buffer was dispensed to reaction wells. 200 μM NaCl was used as a negative control. Data were analyzed using GraphPad Prism version 10 using nonlinear regression analysis (GraphPad Software, La Jolla, CA, USA).

### 4.9. Isothermal titration calorimetry

Experiment was carried out on MicroCal iTC 200 instrument (GE Healthcare). The experiment was performed at 25 °C, 6 μCal/s reference power, and 750 rpm stirring speed. All samples were prepared in 30 mM HEPES, pH 7.5, 150 mM NaCl, 0.5 mM TCEP, and 0.5% (v/v) final DMSO concentration. The samples were degassed at 25 °C right before loading to the sample well and syringe. 50 µM protein was loaded into the sample cell and 500 µM compound was loaded into the syringe. The compound was titrated to the sample cell for 17 cycles. The first injection was with 0.4 µL of titrant, followed by 16 injections with 2.4 µL for each and 180 s intervals. The data were analyzed using Origin 7.0 (OriginLab).

### 4.10. ADP-ribosyl hydrolase activity assay

We used MARylated SRPK2 which was modified at glutamate and aspartate by PARP10 for a substrate. Nanoluciferase-fused ADP-ribosyl superbinder eAf1521 (Nluc-eAf1521) was used as a luminescent probe to detect the remaining MARylated SRPK2 after the hydrolysis [53]. MARylated SRPK2 was prepared as described earlier [23]. To test the hydrolysis activity, 100 nM CHIKV macrodomain was incubated with 0.5 µM MARylated SRPK2 in the presence and absence of tested compounds at room temperature (25 °C) for 2 h. After incubation time, 1 μL of the reaction solution was manually spotted onto the nitrocellulose membrane. After the spots completely dryed out, the membrane was blocked with 5% (w/v) skim milk in Tris-buffered saline with 0.1% Tween-20 (TBS-T) on a shaker for 1 h at 4 °C. Then the membrane was incubated with 3.2 μg/mL Nluc-eAf1521 in 1% (w/v) skim milk solution for 1 h. The membrane was shortly rinsed three times and incubated with TBS-T on a shaker for 1 h to remove free Nluc-eAf1521. After that, the membrane was rinsed once more and visualized using 500 μL of 1:1000 NanoGlo substrate (Promega, catalogue number: N1110) diluted in 10 mM sodium phosphate buffer pH 7.0.

### 4.11. Selectivity

A volume of 10 nL or 100 nL of 10 mM **1** stock was transferred to the reaction wells using Echo 650 acoustic liquid dispenser (Beckman Coulter Life Sciences, Indianapolis, IN, USA). 1 µM of CFP-tagged ADP-ribosyl binder/eraser was mixed with 5 µM MARylated YFP-GAP in the assay buffer as described earlier [53]. 200 µM ADP-ribose was used as a negative control. 10 µL FRET mixture was dispensed to the reaction wells using Mantis liquid handler (Formulatrix, Bedford, MA, USA). For SFV, 1 µM of CFP-SFV was mixed with 5 µM PARylated YFP-GAP, 10 µM untagged SFV was used as the negative control.

### 4.12. Crystallization

The sitting drop vapor diffusion method in TTP Q 96-well plate was employed. 20 mg/mL protein was mixed with reservoir solution to make three drop ratios (75 nL:150 nL or 100 nL:100 nL or 150 nL:75 nL) using Mosquito pipetting robot (TTP Labtech). Crystallization condition was 100 mM Tris, pH 8.0, 15% (w/v) polyvinylpyrrolidone K15, 25% (w/v) polyethylene glycol monomethyl ether 5000. The protein stored buffer was 30 mM Hepes, pH 7.5, 300 mM NaCl, 10% (v/v) glycerol, and 0.5 mM TCEP. Protein diltion buffer was 20 mM Hepes, pH 7.5, and 150 mM NaCl.

The crystallization plates were incubated at 20 °C and monitored by Imager and Icebear [73]. Crystals grew within 1 to 3 days. 2 mM compound (final DMSO concentration was 2%) was prepared in the soaking buffer (60 mM Tris, pH 8.0, 10% (w/v) polyvinylpyrrolidone K15, 15% (w/v) polyethylene glycol monomethyl ether 5000, and 40% glycerol). 0.4 μL of 2 mM compound solution was added to the top of the droplets. The droplets were then incubated for 2 h at 20 °C. Crystals were fished, flashed cool in liquid nitrogen and sent to Diamond. X-ray diffraction data were collected on beamline i04, Diamond, UK.

The dataset was processed by XDS program package [74,75] (**Table S4**). The structures were solved by the method of molecular replacement using Phaser [76]. The CHIKV macrodomain structure (PDB: 6VUQ) was used as a search model. Model building was performed using Coot [77] and model refinement was done using REFMAC5 [78] (**Table S4**). Model was validated using Molprobity. The structures were visualized in PyMOL version 1.7.2.1 (Schrödinger).

### 4.13. Cells and viruses

BHK-21 (baby hamster kidney; ATCC CCL-10) and HEK-293T (human embryonic kidney; gift from Sarah Butcher, University of Helsinki) cells were cultured in Dulbecco’s modified essential media supplement with 10% heat-inactivated Fetal bovine serum (Gibco), 2 mM L-glutamine (Gibco), 100 U/mL penicillin (Gibco), and 100 µg/mL streptomycin (Gibco). The cell lines were passaged every 2-3 days and maintained in a humidifying incubator at 37℃ with 5% CO2.

An attenuated strain of CHIKV, 181/25 [64], was used for studying the effect of the inhibitor on alphavirus replication. This strain was further modified by addition of luciferase reporter as described [65] (gift from Andres Merits, University of Tartu) and further referred to as CHIKV-nLuc in this study.

The cDNA of CHIKV-nLuc was linearized with NotI-HF (New England Biolabs) and capped viral RNA was transcribed *in vitro* using mMESSAGE mMachine SP6 transcription kit (Thermo Fisher Scientific). The virus was rescued by transfecting BHK-21 with the generated infectious RNA using Lipofectamine MessengerMax (Thermo Fisher Scientific) using the manufacturer’s protocol. The supernatant containing the virus was collected 24 h post-transfection and the viral stocks were amplified by infecting BHK-21 cells at multiplicity of infection (MOI) 0.01 plaque forming units (PFU) per mL for 18 h. The CHIKV-nLuc titre estimations were done using plaque assay on BHK-21 cells [79].

To study the effect of the inhibitor on CHIKV replication in cell culture conditions, BHK-21 or HEK-293T cells were seeded on a 96-well flatbottom plate. Approximately 24 h later, the cells were infected with CHIKV-nLuc at MOI 0.01 (BHK-21) or MOI 0.05 (HEK-293T) in presence of indicated dilutions of **1**. At 16 h (BHK-21) or 24 h (HEK-293T) post-infection, the replication of CHIKV-nLuc was determined using NanoGlo Luciferase assay (Promega) based on manufacturer’s instructions. In parallel, non-infected cells with the same dilution of the inhibitor were used to estimate potential toxicity using Cell Titre Glo 2.0 (Promega). 0.5 µM Obatoclax (OLX) was used an independent positive control in these experiments [80]. The luminescence measurements were done using Varioskan LUX microplate reader (Thermo Fisher Scientific).

The analyses were done using Microsoft Excel and R (version 4.4.1) using the package multcomp [81] for post-hoc test. OriginPro (2025) was used to plot the graphs. The number of repeats and the statistical test used to determine the significance are indicated in respective figure legends.

### 4.14. Chemistry

Compound library used in screening (SPECS) was acquired through the Finnish Institute of Molecular Medicine (FIMM). Commercially available structural analogs of the hit compound were acquired from MolPort. Commercial reagents for syntheses were acquired from Tokyo Chemical Industries, Thermo Fisher Scientific, and FluoroChem and the reagents were used as received. ^1^H (400 MHz or 500 MHz) and ^13^C (100 MHz or 125 MHz) nuclear magnetic resonance (NMR) spectra were measured with Bruker Avance III 400 or Bruker Avance 500 HD III spectrometer. Samples were dissolved in CDCl_3_ or *d*_6_-DMSO and placed in 5 mm NMR tubes. All coupling constants are reported in Hz. NMR and HPLC data were processed with Spectrus Processor 2019.1.1 program (ACDLabs). ESI+ TOF MS characterizations were performed using Thermo Scientific QExactive Plus Hybrid Quadrupole-Orbitrap high resolution mass spectrometer with ACQUITY UPLC® BEH C18 column, and the compounds were detected in positive mode. Purities of the compounds were assessed with Shimadzu HPLC instrument using water+0.1% TFA (A) and acetonitrile+0.1% TFA (B) as a mobile phase (gradient B 5% to 90% over 10 minutes) with Waters Atlantis T3 column and detection at 220 nm and 260 nm wavelengths with Shimadzu SDP-10AVP UV-vis detector. Purities of all synthesized final compounds were determined to be >95%. All reactions were monitored with thin-layer chromatography using silica gel-coated aluminum sheets.

### 4.15. Synthesis

#### 2-chloro-1-(2,3-dimethyl-1H-indol-1-yl)ethan-1-one (8a)

2,3-Dimethylindole (1.452 g, 10 mmol, 1 equiv.) was dissolved in 20 mL of 4Å molecular sieve-dried and Ar-bubbled toluene in a 2-necked round bottom flask. The solution was brought to a reflux. Chloroacetyl chloride (1.2 mL, 15 mmol, 1.5 equiv.) was mixed with 3 mL of dried and Ar-bubbled toluene and added with a syringe to the refluxing solution in a dropwise fashion over 10 minutes. The reaction was allowed to proceed at reflux for 18.5 hours. Next, the reaction mixture was allowed to cool to room temperature, and the reaction was quenched with an addition of 20 mL of 1% aqueous NaHCO_3_ solution. The mixture was transferred to an extraction funnel and further diluted with 10 mL of toluene. The organic phase was washed with 30 mL of deionized water and 30 mL of brine. The organic phase was further dried with Na_2_SO_4_, filtered, and evaporated to dryness. The evaporation residue was subjected to flash chromatography (SiO_2_, toluene:*n*-hexane 10:1 isocratic elution). The purified product was collected as a beige solid (1.71 g) in 77% yield. ^1^H NMR (400 MHz, CHLOROFORM-*d*): *δ* ppm 2.21 (s, 3H), 2.60 (s, 3H), 4.71 (s, 2H), 7.27–7.35 (m, 2H), 7.42–7.49 (m, 1H), 7.87–7.96 (m, 1H). / ^1^H NMR (400 MHz, DMSO-*d*_6_): *δ* ppm 2.17 (s, 3H), 2.55 (s, 3H), 5.14 (s, 2H), 7.19–7.34 (m, 2H), 7.41–7.57 (m, 1H), 8.04–8.18 (m, 1H).

#### 2-[(4,6-dihydroxypyrimidin-2-yl)sulfanyl]-1-(2,3-dimethyl-1H-indol-1-yl)ethan-1-one (1r)

2-Thiobarbituric acid (144.6 mg, 1 mmol, 1 equiv.) along with sodium hydroxide (40.3 mg, 1 mmol, 1 equiv.) was added in a reaction tube and dissolved in 2 mL of deionized water with stirring. Compound **8a** (221.7 mg, 1 mmol, 1 equiv.) was dissolved in 2 mL of THF, and the mixture was added to the reaction tube. Tetrabutylammonium bromide (TBAB, 16.1 mg, 50 µmol, 5 mol%) was added to the reaction tube, the tube was sealed and transferred to an oil bath (55 °C). The reaction was allowed to proceed for three days, after which the reaction mixture was transferred to a round-bottom flask, rinsing with THF. The THF was evaporated with a rotary evaporator, and 15 mL of deionized water was added. The mixture was brought to reflux, allowed to cool to room temperature and filtered, washing with 3.5 mL of absolute ethanol. The precipitate was transferred to a flask and triturated twice with boiling absolute ethanol (á 5 mL) and filtered into a sintered funnel, washing with absolute ethanol. The product was dried in a vacuum. The reaction product was isolated as a peach-colored powder (185 mg) in 56% yield. ^1^H NMR (500 MHz, DMSO-*d*_6_): *δ* ppm 2.19 (s, 3H), 2.58 (s, 3H), 4.83 (s, 2H), 5.19 (br s, 1H), 7.17–7.33 (m, 2H), 7.44–7.58 (m, 1H), 8.00–8.15 (m, 1H), 11.42 (br s, 1H), 12.42 (br s, 1H). ^13^C NMR (101 MHz, DMSO-*d*_6_): *δ* ppm 8.5, 14.1, 38.0, 85.7, 115.0, 115.4, 118.2, 123.1, 123.9, 130.7, 132.4, 135.2, 163.0, 167.6, 167.8. HRMS (ESI+, TOF) m/z: [M+H]^+^ calcd for C_16_H_15_N_3_O_3_S 330.0906; found 330.0904.

#### 2-chloro-1-(3-methyl-1H-indol-1-yl)ethan-1-one (8b)

3-Methylindole (659 mg, 5 mmol, 1 equiv.) was dissolved in 10 mL of 4 Å molecular sieve-dried and Ar-bubbled toluene in a 2-necked round bottom flask. The solution was brought to a reflux.

Chloroacetyl chloride (0.6 mL, 7.5 mmol, 1.5 equiv.) was mixed with 1.9 mL of dried and Ar-bubbled toluene, and added with a syringe to the refluxing solution in a dropwise fashion over 20 minutes. The reaction was allowed to proceed at reflux for 16 hours. Next, the reaction mixture was allowed to cool to room temperature, diluted with 10 mL of toluene, and quenched by adding 20 mL of 1% aqueous NaHCO_3_ solution. The mixture was transferred to an extraction funnel. The organic phase was washed with 20 mL of 1% aqueous NaHCO_3_ solution, 20 mL of deionized water, and 20 mL of brine. The organic phase was further dried with MgSO_4_, filtered, and evaporated to dryness. The evaporation residue was subjected to flash chromatography (SiO_2_, isocratic elution with toluene). The purified product was collected as an off-white crystalline solid (824 mg) in 79% yield. ^1^H NMR (400 MHz, CHLOROFORM-*d*): *δ* ppm 2.31 (d, *J*=1.34 Hz, 3H), 4.55 (s, 2H), 7.19 (s, 1H), 7.34 (td, *J*=7.58, 1.10 Hz, 1H), 7.40 (td, *J*=7.46, 0.86 Hz, 1H), 7.52 (dd, *J*=7.21, 0.61 Hz, 1H), 8.42 (br d, *J*=7.95 Hz, 1H).

#### 2-[(4,6-dihydroxypyrimidin-2-yl)sulfanyl]-1-(3-methyl-1H-indol-1-yl)ethan-1-one (9)

2-Thiobarbituric acid (288.8 mg, 2 mmol, 1 equiv.) along with sodium hydroxide (79.2 mg, 2 mmol, 1 equiv.) was added in a reaction tube and dissolved in 4 mL of deionized water with stirring. Compound **8b** (414.8 mg, 2 mmol, 1 equiv.) was dissolved in 4 mL of THF, and the mixture was added to the reaction tube. TBAB (32.2 mg, 100 µmol, 5 mol%) was added to the reaction tube, the tube was sealed and transferred to an oil bath (55 °C). The reaction was allowed to proceed for 67 hours after which the reaction mixture was transferred to a round-bottom flask, rinsing with THF. The solvent was evaporated with a rotary evaporator, and 25 mL of deionized water was added. The mixture was brought to reflux, allowed to cool to room temperature and filtered, washing with 20 mL of absolute ethanol and 3 mL of toluene. The product was dried in a vacuum oven. The reaction product was isolated as an off-white powder (540 mg) in 86% yield. ^1^H NMR (500 MHz, DMSO-*d*_6_): *δ* ppm 2.27 (d, *J*=0.92 Hz, 3H), 4.75 (s, 2H), 5.22 (br s, 1H), 7.28–7.38 (m, 2H), 7.56–7.62 (m, 1H), 7.82 (d, *J*=1.07 Hz, 1H), 8.28–8.30 (m, 1H), 11.80 (br s, 2H). ^13^C NMR (101 MHz, DMSO-*d*_6_): *δ* ppm 9.4, 34.7, 85.7, 115.9, 117.6, 119.1, 123.2, 123.6, 125.0, 131.1, 135.4, 162.9, 165.9, 167.7. HRMS (ESI+, TOF) m/z: [M+H]^+^ calcd for C_15_H_13_N_3_O_3_S 316.0750; found 316.0748.

#### 2-chloro-1-(1,2,3,4-tetrahydro-9H-carbazol-9-yl)ethan-1-one (8c)

1,2,3,4-Tetrahydrocarbazole (857 mg, 5 mmol, 1 equiv.) was dissolved in 10 mL of 4Å molecular sieve-dried and Ar-bubbled toluene in a 2-necked round bottom flask. The solution was brought to a reflux. Chloroacetyl chloride (0.6 mL, 7.5 mmol, 1.5 equiv.) was mixed with 1.9 mL of dried and Ar-bubbled toluene, and added with a syringe to the refluxing solution in a dropwise fashion over 10 minutes. The reaction was allowed to proceed at reflux for 19 hours. Next, the reaction mixture was allowed to cool to room temperature, diluted with 10 mL of toluene, and quenched by adding 20 mL of 1% aqueous NaHCO_3_ solution. The mixture was transferred to an extraction funnel. The organic phase was washed with 20 mL of 1% aqueous NaHCO_3_ solution, 20 mL of deionized water, and 20 mL of brine. The organic phase was further dried with MgSO_4_, filtered, and evaporated to dryness. The evaporation residue was subjected to Flash chromatography (SiO_2_, isocratic elution with toluene). The purified product was collected as a yellowish solid (1.11 g) in 90% yield. ^1^H NMR (400 MHz, CHLOROFORM-*d*): *δ* ppm 1.79–1.97 (m, 4H), 2.59–2.72 (m, 2H), 2.95–3.05 (m, 2H), 4.64 (s, 2H), 7.23–7.33 (m, 2H), 7.37–7.44 (m, 1H), 7.97–8.08 (m, 1H).

#### 2-[(4,6-dihydroxypyrimidin-2-yl)sulfanyl]-1-(1,2,3,4-tetrahydro-9H-carbazol-9-yl)ethan-1-one (10)

2-Thiobarbituric acid (289.8 mg, 2 mmol, 1 equiv.) along with sodium hydroxide (81.5 mg, 2 mmol, 1 equiv.) was added in a reaction tube and dissolved in 4 mL of deionized water with stirring. Compound **8c** (497.2 mg, 2 mmol, 1 equiv.) was dissolved in 4 mL of THF, and the mixture was added to the reaction tube. TBAB (32.7 mg, 100 µmol, 5 mol%) was added to the reaction tube, the tube was sealed and transferred to an oil bath (55 °C). The reaction was allowed to proceed for 69 hours after which the solvent system was decanted away, and solids were transferred to a round-bottom flask, rinsing with THF, and evaporated to dryness. The crude product was triturated three times with 10 mL of cold toluene. To the crude product, 15 mL of deionized water was added. The mixture was brought to reflux, allowed to cool to room temperature, and filtered, washing with 10 mL of absolute ethanol. Next, the crude product was triturated with a total of 30 mL of methanol. The product was dried in a vacuum oven. The reaction product was isolated as a pale pink powder (562 mg) in 79% yield. ^1^H NMR (400 MHz, DMSO-*d*_6_): *δ* ppm 1.73–1.95 (m, 4H), 2.64 (br s, 2H), 3.05 (br s, 2H), 4.79 (s, 2H), 5.19 (br s, 1H), 7.21–7.34 (m, 2H), 7.40–7.54 (m, 1H), 8.08–8.20 (m, 1H), 11.41 (br s, 1H), 12.37 (br s, 1H). ^13^C NMR (101 MHz, DMSO-*d*_6_): *δ* ppm 20.7, 21.4, 23.4, 25.8, 37.6, 85.6, 115.7, 117.7, 117.8, 123.2, 124.1, 129.7, 135.0, 135.5, 163.0, 167.4, 167.6. HRMS (ESI+, TOF) m/z: [M+H]^+^ calcd for C_18_H_17_N_3_O_3_S 356.1063; found 356.1053.

#### 2-chloro-1-(1,2-dimethyl-3H-benzo[e]indol-3-yl)ethan-1-one (8d)

1,2-Dimethyl-3H-benz[e]indole (587 mg, 3 mmol, 1 equiv.) was dissolved in 6 mL of 4Å molecular sieve-dried and Ar-bubbled toluene in a 2-necked round bottom flask. The solution was brought to a reflux. Chloroacetyl chloride (360 µL, 4.5 mmol, 1.5 equiv.) was mixed with 1.2 mL of dried and Ar-bubbled toluene and added with a syringe to the refluxing solution in a dropwise fashion over 10 minutes. The reaction was allowed to proceed at reflux for 22 hours. Next, the reaction mixture was allowed to cool to room temperature, diluted with 10 mL of toluene, and quenched by adding 15 mL of 1% aqueous NaHCO_3_ solution. The mixture was transferred to an extraction funnel. The organic phase was washed with 20 mL of deionized water and 20 mL of brine. The organic phase was filtered through a thin pad of silica gel, rinsed with toluene, and evaporated to dryness. The evaporation residue was subjected to consecutive flash chromatographic separations (SiO_2_, isocratic elution with toluene). The purified product was collected as a beige solid (426 mg) in 52% yield. Note: the compound is unstable in ambient conditions, forming a dark green polymerization product. ^1^H NMR (400 MHz, CHLOROFORM-*d*): *δ* ppm 2.65 (s, 3H), 2.66 (s, 3H), 4.75 (s, 2H), 7.46–7.53 (m, 1H), 7.58 (s, 1H), 7.72 (d, *J*=9.05 Hz, 1H), 7.94 (d, *J*=7.95 Hz, 1H), 8.11 (d, *J*=9.05 Hz, 1H), 8.55 (d, *J*=8.44 Hz, 1H).

#### 2-[(4,6-dihydroxypyrimidin-2-yl)sulfanyl]-1-(1,2-dimethyl-3H-benzo[e]indol-3-yl)ethan-1-one (11)

2-Thiobarbituric acid (108.4 mg, 0.74 mmol, 1 equiv.) along with sodium hydroxide (31.1 mg, 0.74 mmol, 1 equiv.) was added in a reaction tube and dissolved in 2 mL of deionized water with stirring. Compound **8d** (199.7 mg, 0.73 mmol, 1 equiv.) was dissolved in 4 mL of THF, and the mixture was added to the reaction tube. TBAB (12.3 mg, 37 µmol, 5 mol%) was added to the reaction tube, the tube was sealed and transferred to an oil bath (50 °C). The reaction was allowed to proceed for 69 hours after which the mixture was transferred to a round-bottom flask, rinsing with THF, and the mixture was concentrated with rotary evaporator. The crude product was triturated with a total of 20 mL of cold deionized water and 20 mL of cold toluene. The filtered crude product was washed with 7 mL of dichloromethane and a total of 17 mL of *n*-hexane. Next, the precipitate was dissolved in 3 mL of DMSO and heated to 60 °C. The dropwise addition of deionized water (7 mL in total) to the warm solution initiated a precipitation of the reaction product. The mixture was allowed to cool to room temperature and filtered, washing with 30 mL of deionized water and 2 mL of H_2_O/EtOH 1:1-mixture. The pure product was dried in a vacuum oven. The reaction product was isolated as a pink powder (137.2 mg) in 49% yield. ^1^H NMR (400 MHz, DMSO-*d*_6_): *δ* ppm 2.63 (s, 3H), 2.65 (s, 3H), 4.88 (s, 2H), 5.18 (br s, 1H), 7.50 (t, *J*=7.58 Hz, 1H), 7.60 (t, *J*=7.46 Hz, 1H), 7.73 (d, *J*=9.17 Hz, 1H), 7.99 (d, *J*=7.95 Hz, 1H), 8.24 (d, *J*=9.17 Hz, 1H), 8.54 (d, *J*=8.44 Hz, 1H), 11.37 (br s, 1H), 12.49 (br s, 1H). ^13^C NMR (101 MHz, DMSO-*d*_6_): *δ* ppm 12.9, 13.3, 38.2, 85.6, 115.3, 115.9, 123.45, 123.48, 124.0, 124.2, 126.1, 127.5, 128.5, 130.4, 131.1, 132.1, 167.4, 167.7, 169.0. HRMS (ESI+, TOF) m/z: [M+H]^+^ calcd for C_20_H_17_N_3_O_3_S 380.1063; found 380.1076.

#### 2-[(4,6-diethoxypyrimidin-2-yl)sulfanyl]-1-(2,3-dimethyl-1H-indol-1-yl)ethan-1-one (12)

Compound **1r** (131.6 mg, 0.4 mmol, 1 equiv.) along with potassium carbonate (224 mg, 1.6 mmol, 4 equiv.) were weighed into a reaction tube, which was sealed and flushed with a stream of argon gas. Diethyl sulphate (390 µL, 3 mmol, 7.5 equiv.) was added dropwise to the mixture through the septum with simultaneous gentle stirring. The formed slurry was heated with an oil bath (80 °C) for 4 hours. Next, the reaction mixture was dissolved in 4 mL of ethyl acetate, 10 mL of saturated NaHCO_3_ was added, and the mixture was stirred at rt. The mixture was transferred into an extraction funnel and further diluted with 5 mL of deionized water and 5 mL of ethyl acetate. The organic phase was separated, and washed with 10 mL of saturated NaHCO_3_, 10 mL of deionized water, and 10 mL of brine. The organic phase was dried with MgSO_4_, filtered, and evaporated to dryness. The evaporation residue was subjected to flash chromatography (SiO_2_, CHCl_3_ 100 % to CHCl_3_:MeOH 95:5 gradient elution). The pure product was collected as a white solid (78.1 mg) in 51% yield. ^1^H NMR (500 MHz, CHLOROFORM-*d*): *δ* ppm 1.18 (t, *J*=7.10 Hz, 6H), 2.22 (s, 3H), 2.60 (s, 3H), 4.08 (q, *J*=7.17 Hz, 4H), 4.61 (s, 2H), 5.66 (s, 1H), 7.22–7.32 (m, 2H), 7.40–7.50 (m, 1H), 7.91–8.04 (m, 1H). ^13^C NMR (126 MHz, CHLOROFORM-*d*): *δ* ppm 8.7, 14.3, 14.4, 38.9, 62.5, 86.5, 114.9, 115.9, 118.3, 123.1, 123.9, 131.3, 132.7, 135.4, 168.4, 168.6, 170.6. HRMS (ESI+, TOF) m/z: [M+H]^+^ calcd for C_20_H_23_N_3_O_3_S 386.1533; found 386.1521.

#### 2-[(5-bromo-4,6-dihydroxypyrimidin-2-yl)sulfanyl]-1-(2,3-dimethyl-1H-indol-1-yl)ethan-1-one (13)

Compound **1r** (201.1 mg, 0.61 mmol, 1 equiv.) was weighed into a reaction tube, and 1.2 mL of DMSO was added with stirring. After the starting material dissolved completely, *N*-bromosuccinimide (119.3 mg, 0.67 mmol, 1.1 equiv.) was added in one portion to the reaction tube. The reaction was allowed to proceed with stirring for 2.5 hours, after which it was quenched with 10 mL of deionized water. The formed precipitate was filtered into a sintered funnel, washed with ca. 70 mL of deionized water, 30 mL of 1:1 mixture of water and ethanol, and 5 mL of ethanol. The precipitate was dried in a vacuum oven (55 °C). The pure product was collected as a peach-colored solid (199 mg) in 80% yield. ^1^H NMR (400 MHz, DMSO-*d*_6_): *δ* ppm 2.19 (s, 3H), 2.59 (s, 3H), 4.89 (s, 2H), 7.20–7.37 (m, 2H), 7.46–7.56 (m, 1H), 8.05–8.18 (m, 1H), 12.62 (br s, 2H). ^13^C NMR (101 MHz, DMSO-*d*_6_): *δ* ppm 8.5, 14.2, 38.5, 83.1, 115.2, 115.5, 118.2, 123.2, 124.0, 130.7, 132.4, 135.3, 159.6, 163.6, 167.2. HRMS (ESI+, TOF) m/z: [M+H]^+^ calcd for C_16_H_14_BrN_3_O_3_S 408.0012; found 408.0025.

## Acknowledgments

We thank Dr. Johan Pääkkönen and Dr. Anil Sohail for their discussion in crystallography and helping with the data collection. The use of the facilities and expertise of the Biocenter Oulu Structural Biology core facility (a member of Biocenter Finland, Instruct-ERIC Centre Finland and FINStruct), Proteomics and Protein Analysis core facility (a member of Biocenter Finland) and Biocenter Oulu sequencing center are gratefully acknowledged. We are grateful to local contacts at the Diamond for assistance in using beamline i04.

## CRediT authorship contribution statement

**Men Thi Hoai Duong**: Methodology, Investigation, Data curation, Formal analysis, Visualization, Writing-original draft, Writing-review and editing; **Tomi A.O. Parviainen**: Methodology, Investigation, Data curation, Formal analysis, Visualization, Writing-original draft, Writing review and editing; **Aditya Thiruvaiyaru**: Methodology, Investigation, Data curation, Formal analysis, Visualization, Writing-original draft, Writing-review and editing; **Tero Ahola**: Writing-review and editing, Supervision; **Juha P. Heiskanen**: Writing-review and editing, Supervision; **Lari Lehtiö**: Conceptualization, Validation, Writing-review and editing, Supervision.

## Data availability

Atomic coordinates and structure factors will be available at the Protein Data Bank with the ID 9TDQ. Raw diffraction data will be available at fairdata.fi (https://doi.org/10.23729/fd-4324791a-9a90-369e-9e23-cb457d0a3f48). Other study data are included in the article and supporting information.

