## Supplemental Information for "Discovery of dual thiobarbiturate-indole scaffold as a selective inhibitor targeting chikungunya virus nsP3 macrodomain through a cryptic binding pocket"

### Content

**Figure S1.** Multiple sequence alignment.

**Figure S2.** Protein concentration optimization.

**Figure S3.** *Z*'-factors of high-throughput screening and hit validation.

**Figure S4**. Different behavior of Arg1477 in the ADP-ribose complex structure and the compound **1** complex structure.

**Figure S5.** $IC_{50}$ of compound **1** against CHIKV macrodomain R1477F and R1477A mutants.

**Figure S6.** Toxicity assay of compound **1** in cells.

**Table S1.** Static parameters for assay validation.

**Table S2.** 12 compounds stabilizing the CHIKV macrodomain without showing inhibitory activity in the counter screen.

**Table S3**. Physicochemical properties of compound **1**.

**Table S4.** Data collection and model refinement.

**Figure S7-S23.** NMR spectra of synthesized compounds

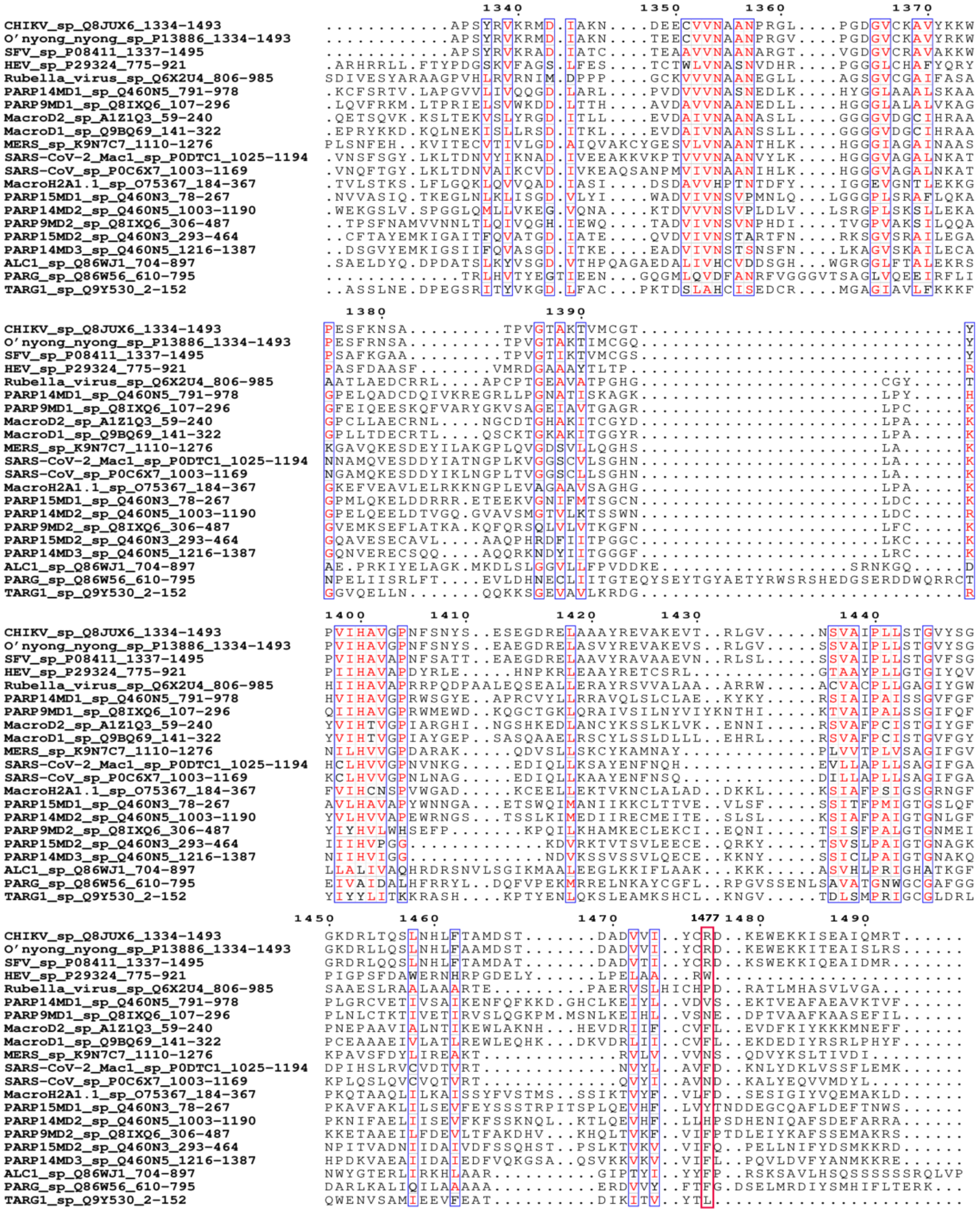

**Figure S1**. **Multiple sequence alignment of human and representative viral macrodomains**.

Residues are numbered according to the CHIKV sequence. The residues in each column are colored based on a similarity score assigned to that column. Columns with scores higher than 0.7

are outlined in blue, and residues are highlighted in red. Arg1477 in the CHIKV macrodomain and corresponding residues in other macrodomains are framed in a red box.

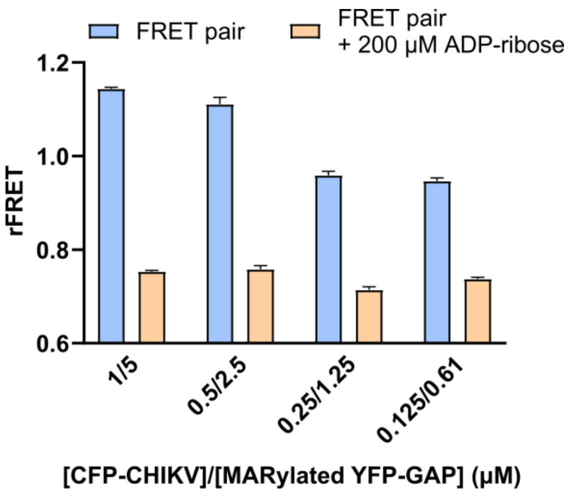


**Figure S2**. **Protein concentration optimization for FRET-based assay**.

The ratio between CFP-CHIKV and MARylated YFP-GAP was kept as 1:5. Data are shown as the mean ± standard deviation (n = 4).

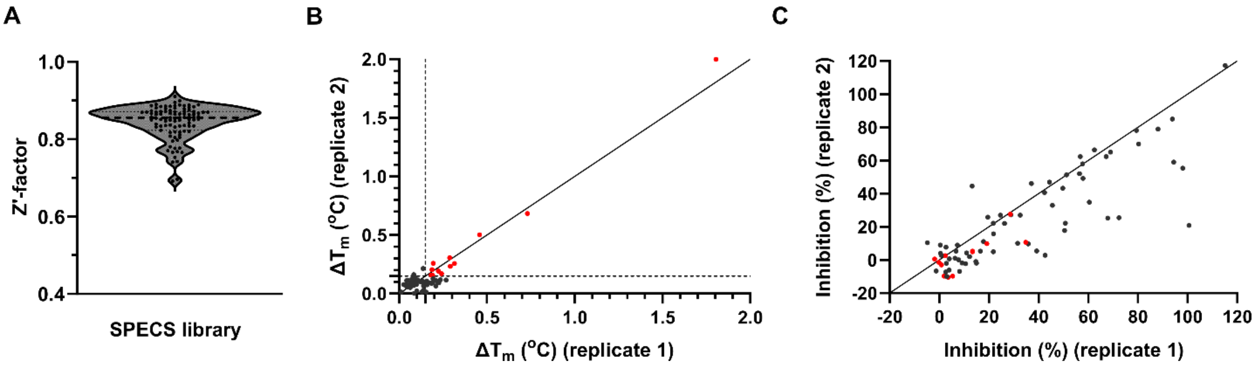


**Figure S3. Z'-factors of high-throughput screening and hit validation.**

**A**) The screening window coefficient Z'-factors across 94 screening plates. Z'-factors are shown as single dots. The mean is shown as a dash line.

**B**) Melting temperature shift of the CHIKV macrodomain in the presence of 77 hit compounds, 30 µM each compound. Data from two replicates are shown. 12 compounds that induced a shift greater than 0.15 °C are highlighted as red dots. The dash lines mark the cut-off at the $\Delta T_m$ of 0.15 $^oC$.

**C**) Counter screen panel of 30 µM of 77 primary hits using the FRET-based assay of CFP-ARC4 and YFP-TBM. Data of two replicates are shown. Among the 12 compounds that induced a thermal shift greater than 0.15 °C, all exhibited less than 40% inhibition in the counter screen, with 10 of these compounds showing less than 20% inhibition. These 12 compounds are marked as red dots.

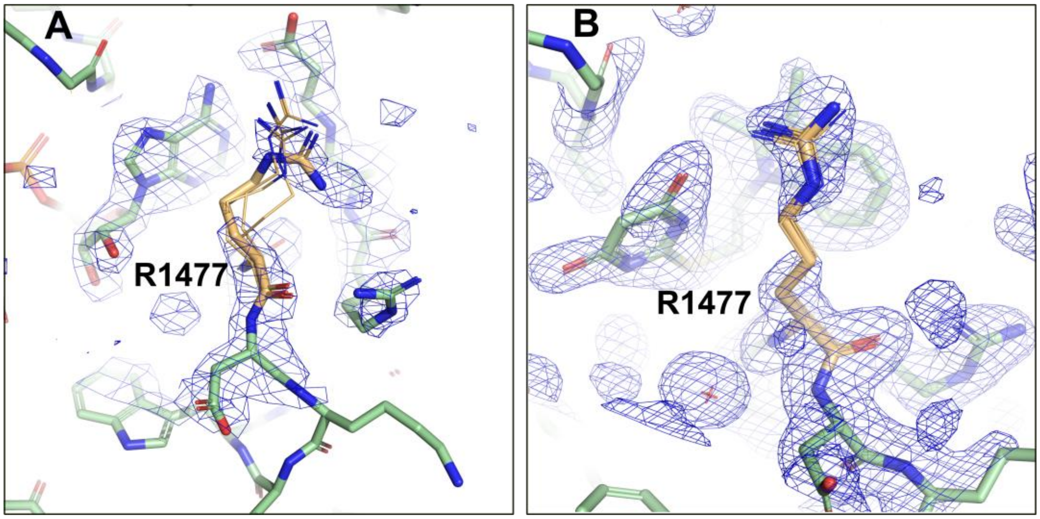


**Figure S4. Different behavior of Arg1477 in the ADP-ribose complex structure and the compound 1 complex structure.**

**A)** The Arg1477 in the ADP-ribose complex (PDB: 3GPO) has multiple conformations and its sidechain has a poorly defined electron density. **B)** The Arg1477 in the compound **1** complex (PDB: 9TDQ) has clearly defined electron density for the guanidinium group stacking with the compound. The 2Fo-Fc maps within 5 Å from the Arg1477 residue are contoured at 0.8 σ and colored in blue.

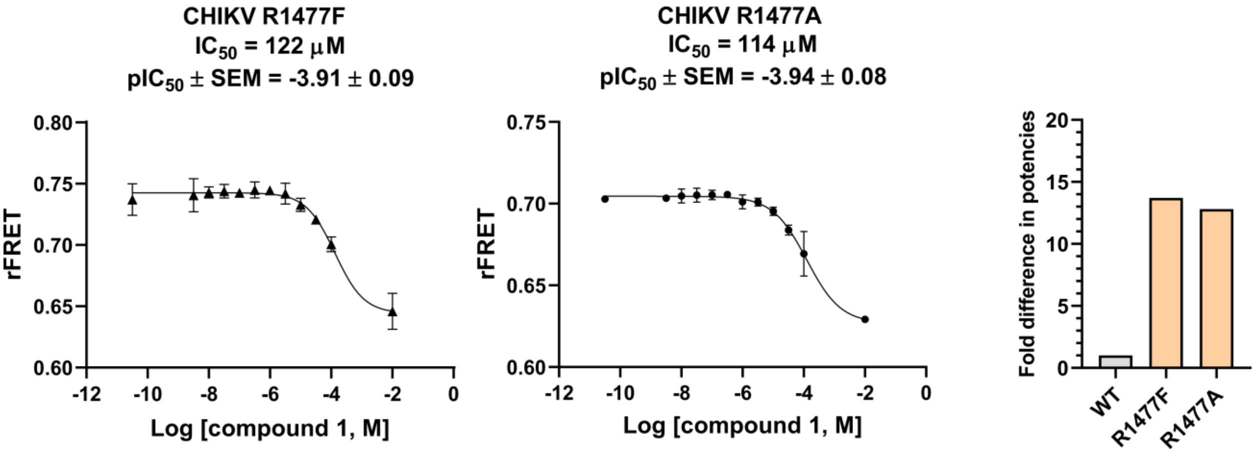


**Figure S5. $IC_{50}$ curves of compound 1 for the CHIKV macrodomain R1477F and R1477A mutants.** The data are presented as mean ± standard deviation from four technical replicates, and $pIC_{50}$ is calculated from three independent experiments.

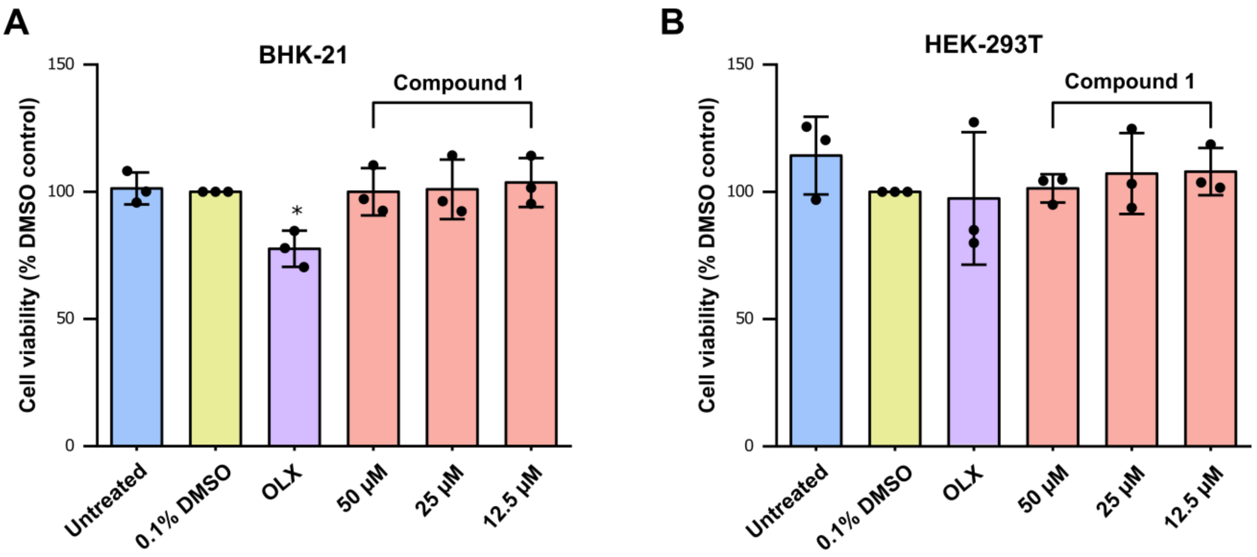


**Figure S6. Toxicity of compound 1**. The cell lines **A)** BHK-21 or **B)** HEK-293T were incubated with the inhibitor and the ATP was quantified at same timepoint as **Figure 6**. The cell viability (based on the ATP) was normalized with respect to the DMSO control and represented as a mean of three independent biological repeats (each carried out in quadruplicate) ± standard deviation. The statistical significance was calculated using one-way ANOVA with Dunnett's multiple comparisons test of each condition against the DMSO control; *$p$<0.05.

**Table S1**. Static parameters of assay validation

| Parameters | Day 1 | Day 2 | Day 3 | | |
|---|---|---|---|---|---|
| | | | Plate 1 | Plate 2 | Plate 3 |
| Positive control AVR (rFRET) ± SD (CV%) | 0.81 ± 0.01 (1.8 %) | 0.81 ± 0.02 (1.9 %) | 0.8 ± 0.01 (1.7 %) | 0.86 ± 0.01 (1.3 %) | 0.84 ± 0.01 (1.4 %) |
| Negative control AVR (rFRET) ± SD (CV %) | 0.57 ± 0.01 (1.3 %) | 0.57 ± 0.01 (1.5 %) | 0.58 ± 0.01 (1.6 %) | 0.63 ± 0.01 (1.3 %) | 0.61 ± 0.01 (1.1 %) |
| Z’-factor (CV%) | 0.72 | 0.71 | 0.69 | 0.76 | 0.76 |
| | 0.73 ± 0.03 (4.4%) | | | | |
| S/B (CV%) | 1.42 | 1.43 | 1.38 | 1.37 | 1.37 |
| | 1.39 ± 0.03 (2.2 %) | | | | |
| S/N (CV%) | 14.65 | 13.80 | 13.29 | 17.17 | 17.03 |
| | 15.19 ± 1.81 (11.9 %) | | | | |
| Plate-to-plate CV (%) Calculated from Z’-factor | 5.7 % | | | | |
| Day-to-day CV (%) Calculated from Z’-factor | 3.4 % | | | | |

**Table S2**. 12 compounds stabilizing CHIKV macrodomain without showing inhibitory activity in the counter screen.

| No. | Structure | % Inhibition (FRET-based assay for the CHIKV macrodomain) | $\Delta T_m$ (ºC) |
|---|---|---|---|
| **1** | 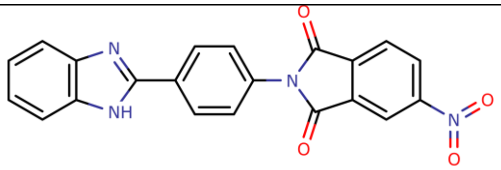 | 50 | 1.90 ± 0.14 |
| **2** | 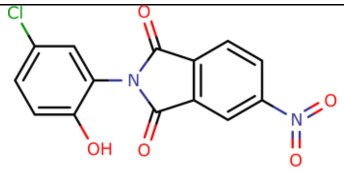 | 65 | 0.71 ± 0.03 |
| **3** | 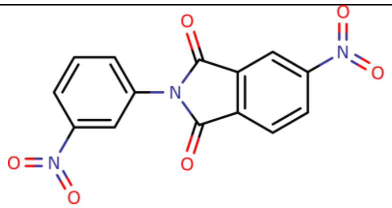 | 48 | 0.48 ± 0.03 |
| **4** | 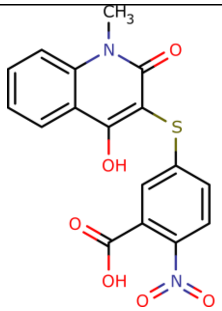 | 28 | 0.30 ± 0.01 |
| **5** | 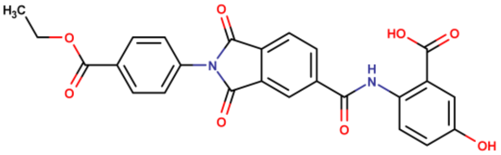 | 20 | 0.29 ± 0.04 |
| **6** | 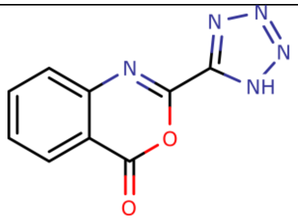 | 22 | 0.26 ± 0.04 |

| | | | |
|---|---|---|---|
| **7** | 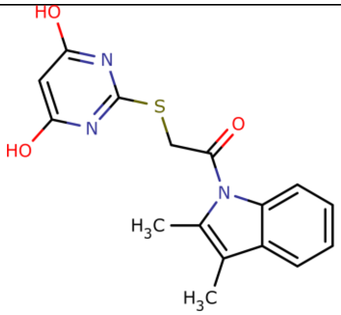 | 27 | 0.23 ± 0.05 |
| **8** | 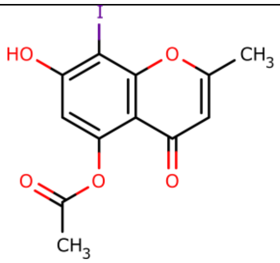 | 42 | 0.21 ± 0.02 |
| **9** | 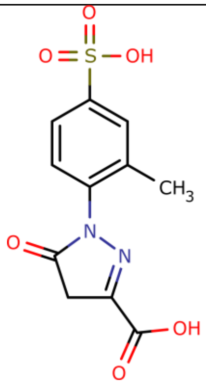 | 26 | 0.21 ± 0.05 |
| **10** | 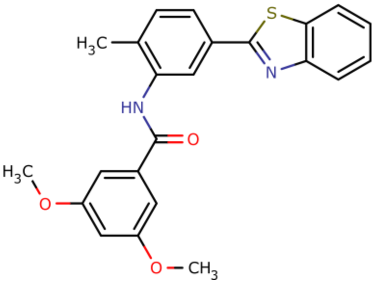 | 33 | 0.17 ± 0.02 |

| | | | |
|---|---|---|---|
| **11** | 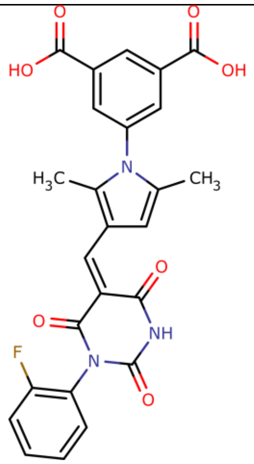 | 43 | 0.17 ± 0.02 |
| **12** | 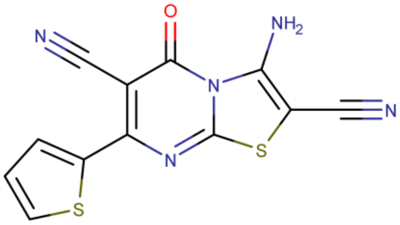 | 22 | 0.20 ± 0.01 |

**Table S3.** Physicochemical properties of compound 1.

| Parameters | Rule | **1** |
| --- | --- | --- |
| MW | < 500 Da | 329.37 Da |
| Number of H-bond acceptors | < 10 | 5 |
| Number of H-bond donors | < 5 | 2 |
| Number of rotatable bonds | < 10 | 4 |
| TPSA (topological polar surface area) | < 140 $Å^2$ | 113.54 $Å^2$ |
| cLogD (pH 7.4) | - | 3.23 |

**Table S4**. Data collection and model refinement statistics for the crystal structure of the CHIKV macrodomain soaked with compound **1**.

| **PDB id.** | **9TDQ** |
|---|---|
| **Data collection** | |
| Beamline | Diamond I04 |
| Wavelength (Å) | 0.953731 |
| Space group | $P3_1$ |
| Unit cell dimensions | |
| *a*, *b*, *c* (Å) | 86.90 86.90 84.96 |
| α, β, γ (°) | 90 90 120 |
| Resolution range (Å) | 50 - 1.65 (1.69 - 1.65) |
| Total no. of reflections | 917278 (68133) |
| No. of unique reflections | 86300 (6448) |
| Completeness (%) | 99.9% (99.9%) |
| $I/\sigma(I)$ | 21.13 (2.16) |
| $CC_{1/2}$ (%) | 100 (74.8) |
| $R_{meas}$ (%) | 5.9 (105.1) |
| **Model building and refinement** | |
| $R_{work}$ | 0.1443 |
| $R_{free}$ | 0.1869 |
| No. of atoms | |
| Protein | 5075 |
| Ligands | 92 |
| Water | 301 |
| RMSD | |
| Bonds (Å) | 0.006 |
| Angles (°) | 1.276 |
| Average *B* factors ($Å^2$) | |
| Protein | 36.00 |
| Ligands | 68.52 |
| Water | 41.46 |
| Ramachandran plot | |
| Favoured (%) | 100 |
| Allowed (%) | 0 |
| Outliers (%) | 0 |

**Figure S7.** $^{1}$H NMR spectrum of compound **8a** in $CDCl_3$ (-0.5 – 15 ppm).

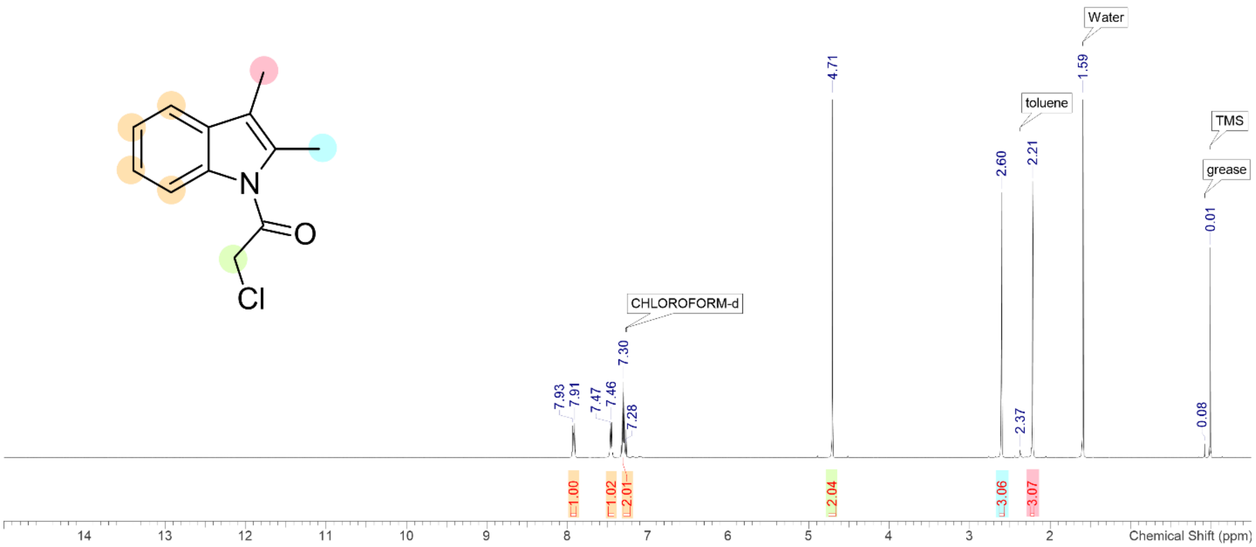


**Figure S8.** $^{1}$H NMR spectrum of compound **1r** in $d_6$-DMSO (-0.5 – 15 ppm).

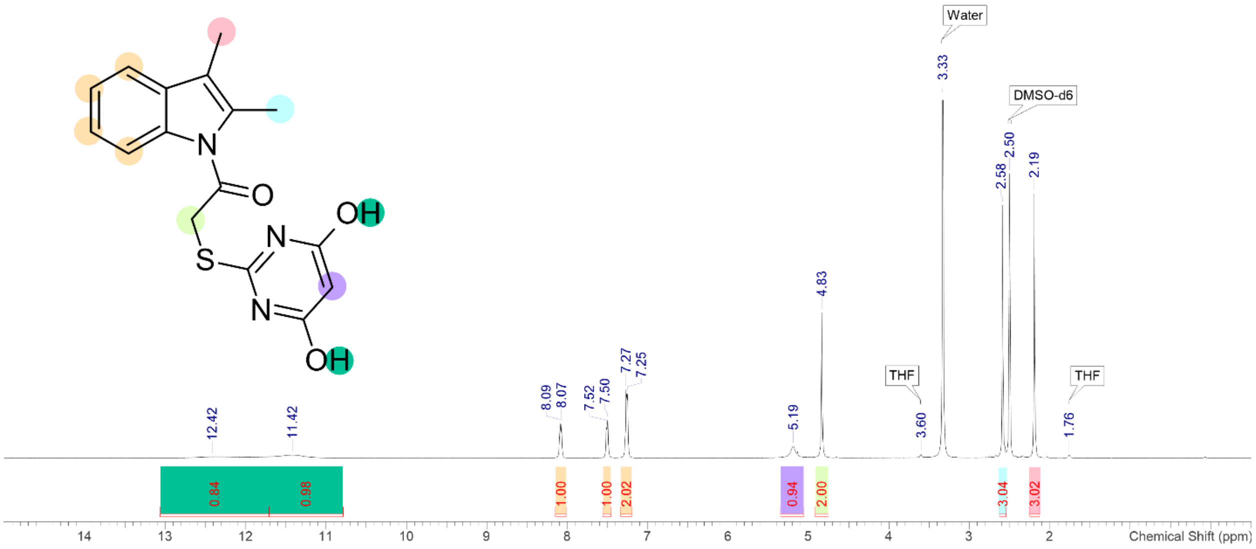

**Figure S9.** $^{13}$C NMR spectrum of compound **1r** in $d_6$-DMSO (0 – 220 ppm).

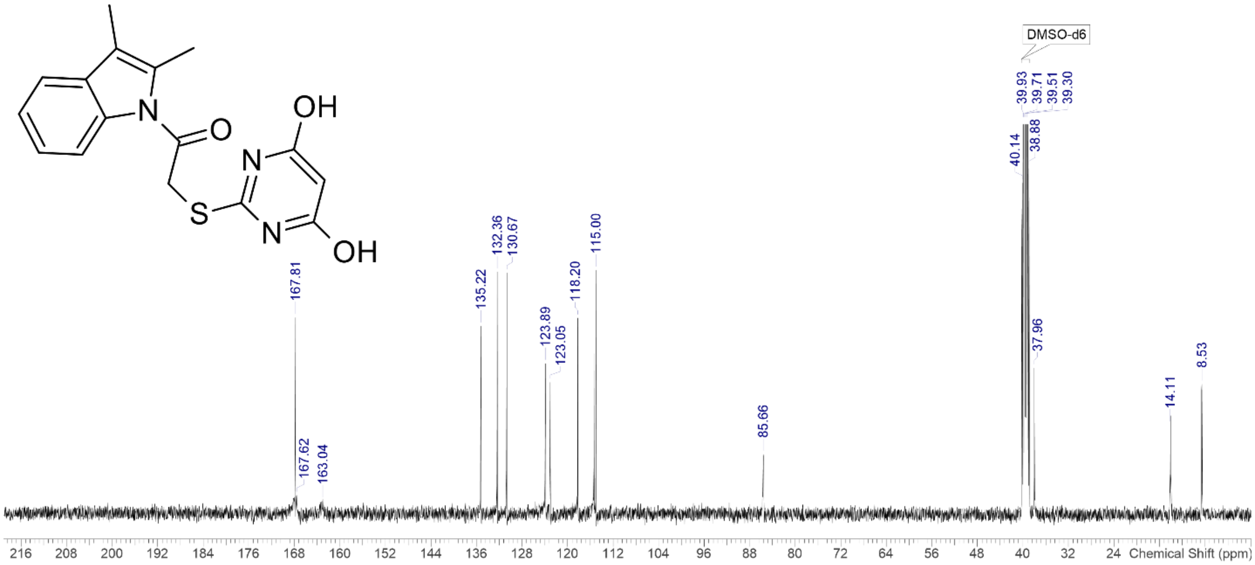


**Figure S10.** $^{1}$H NMR spectrum of compound **8b** in $CDCl_3$ (-0.5 – 15 ppm).

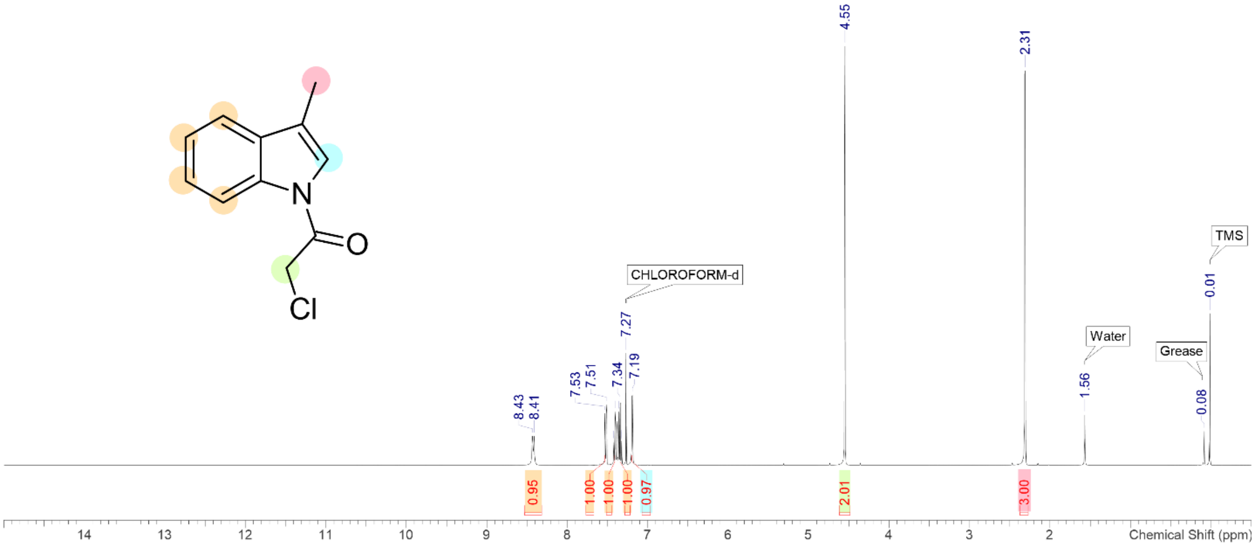

**Figure S11.** $^{1}$H NMR spectrum of compound **9** in $d_6$-DMSO (-0.5 – 15 ppm).

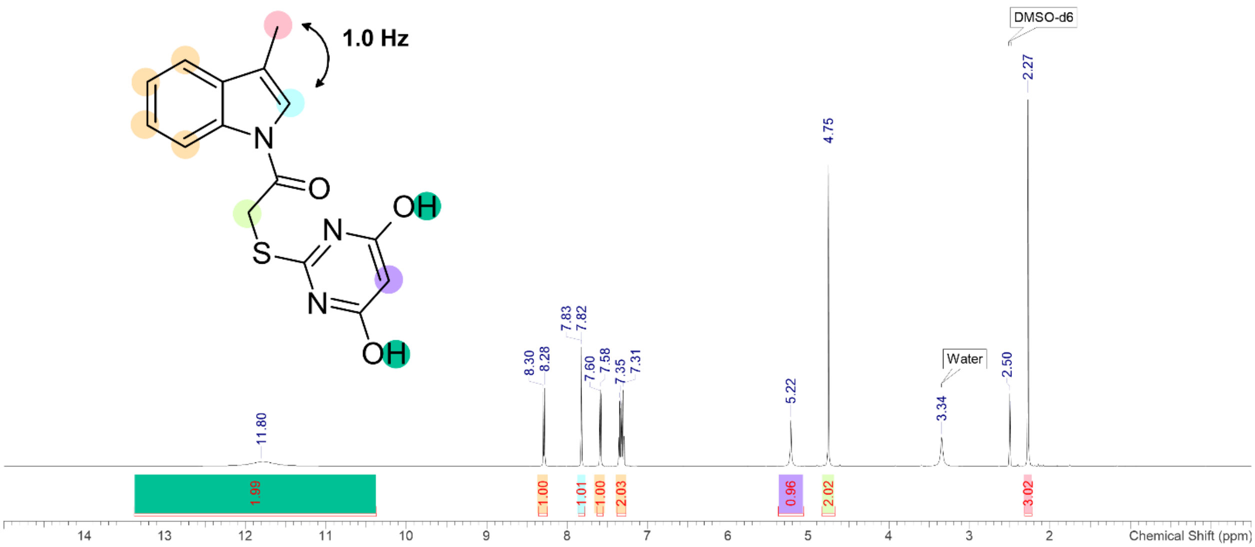


**Figure S12.** $^{13}$C NMR spectrum of compound **9** in $d_6$-DMSO (0 – 220 ppm).

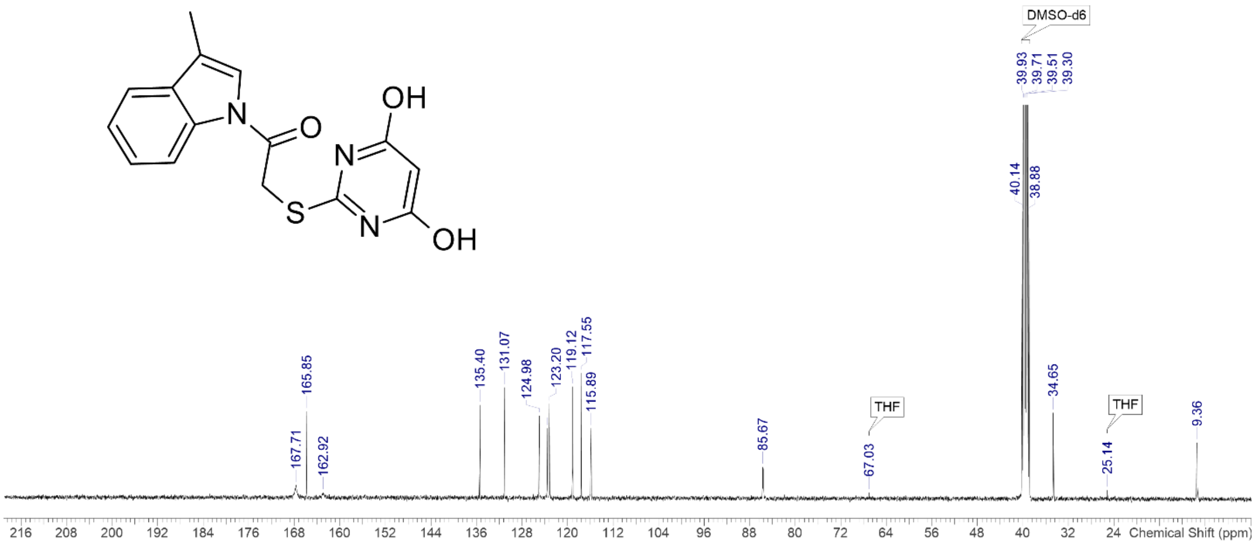

**Figure S13.** [1]H NMR spectrum of compound **8c** in $CDCl_3$ (-0.5 – 15 ppm).

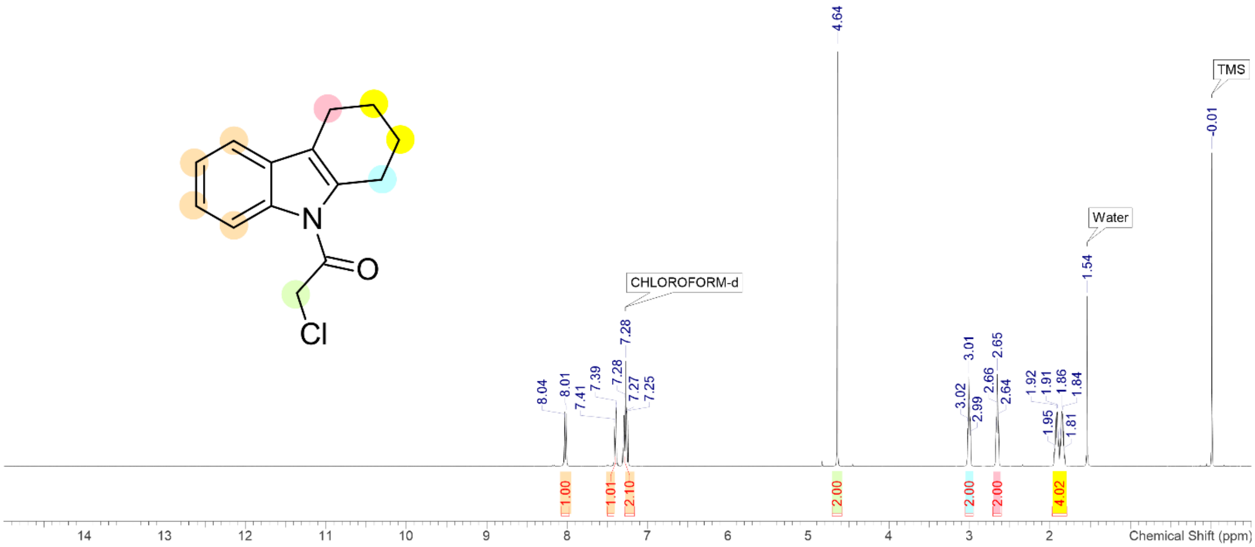


**Figure S14.** [1]H NMR spectrum of compound **10** in $d_6$-DMSO (-0.5 – 15 ppm).

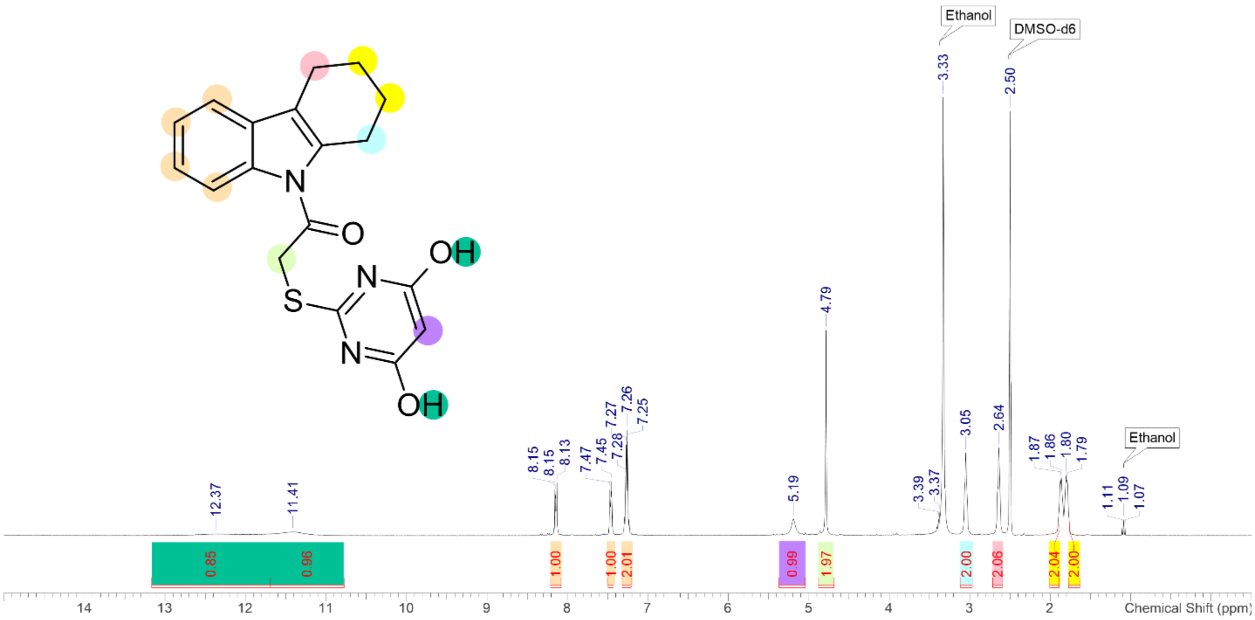

**Figure S15.** $^{13}$C NMR spectrum of compound **10** in $d_6$-DMSO (0 – 220 ppm).

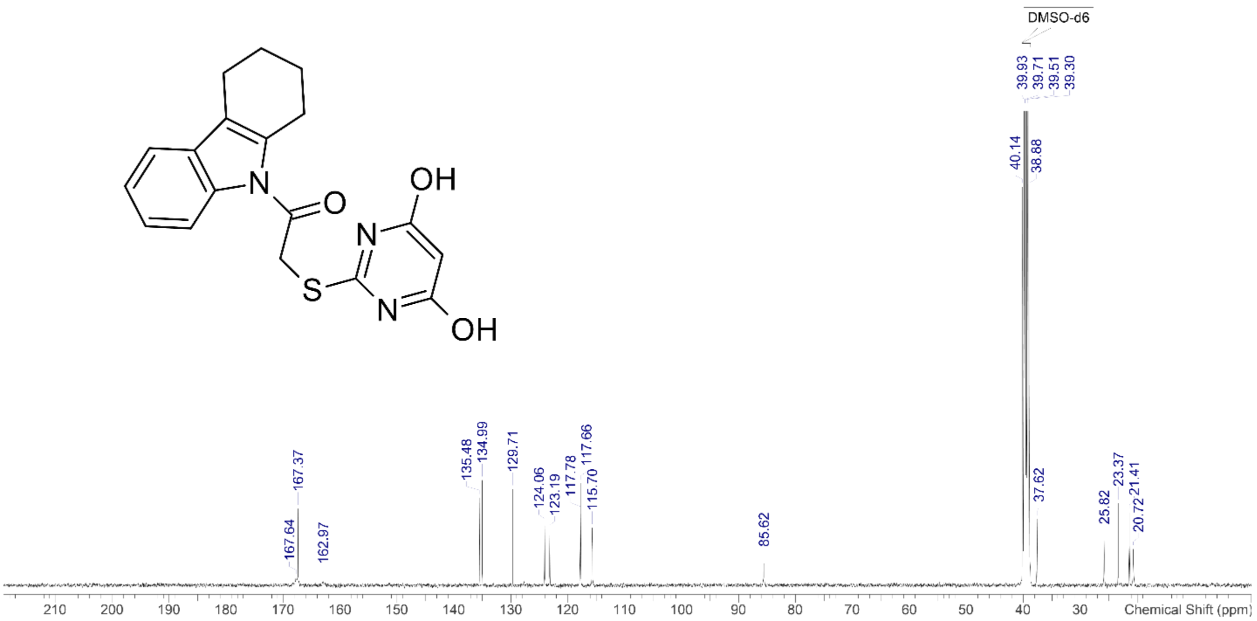


**Figure S16.** $^1$H NMR spectrum of compound **8d** in $CDCl_3$ (-0.5 – 15 ppm).

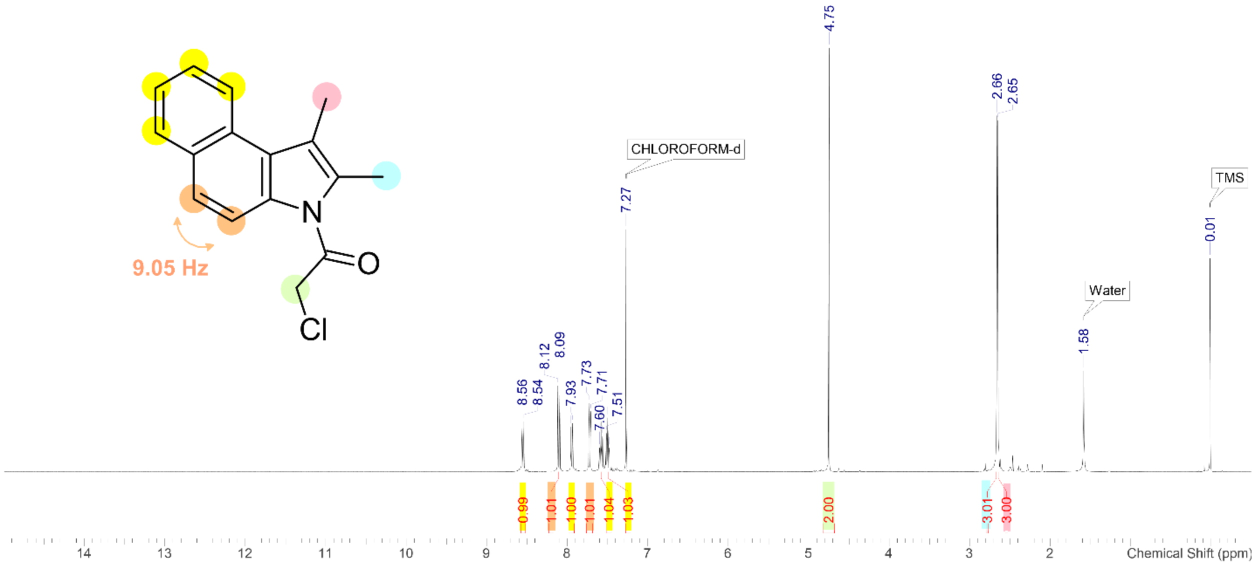

**Figure S17.** $^{1}$H NMR spectrum of compound **11** in $d_6$-DMSO (-0.5 – 15 ppm).

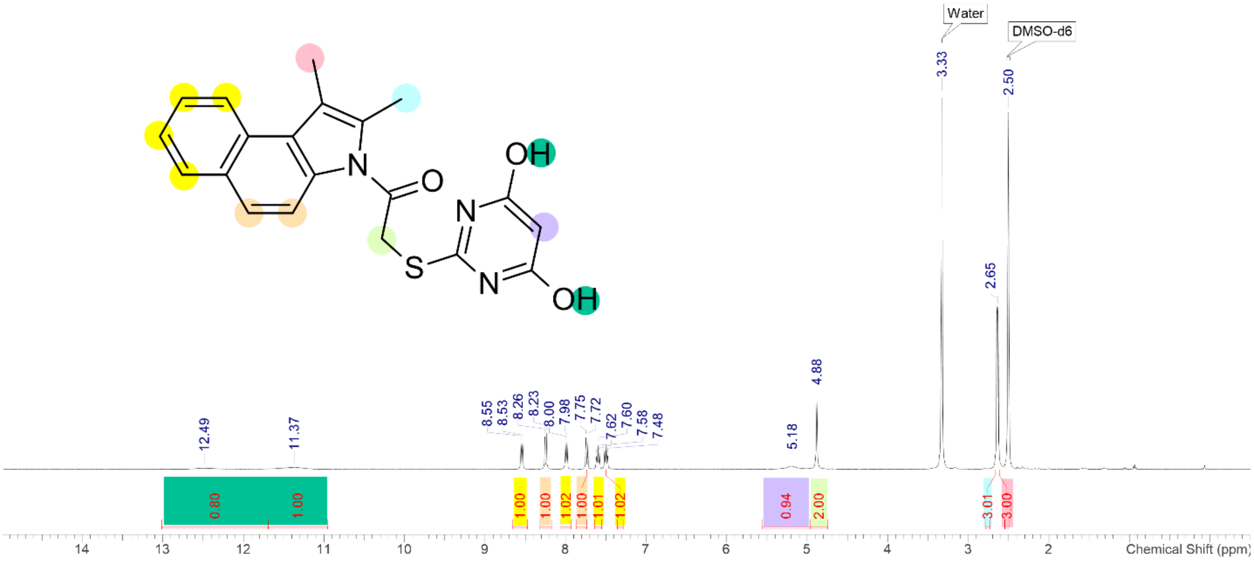


**Figure S18.** Local zoom of $^{1}$H NMR spectrum of compound **11** (7 – 9 ppm).

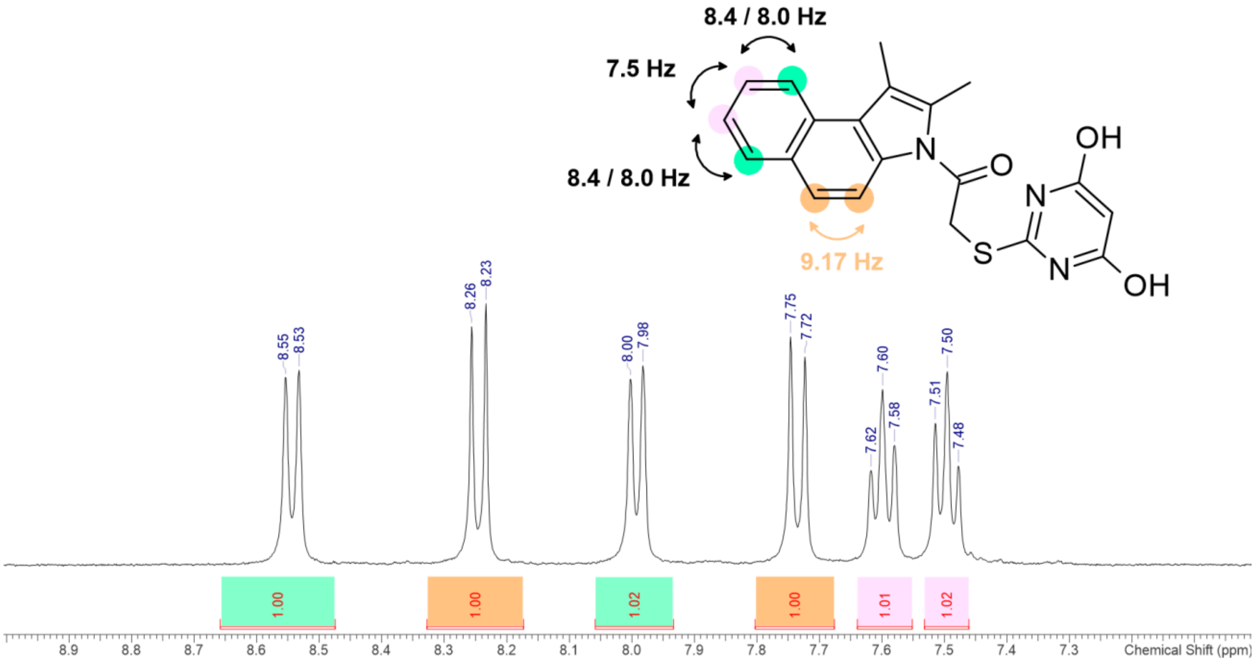

**Figure S19.** $^{13}$C NMR spectrum of compound **11** in $d_6$-DMSO (0 – 220 ppm).

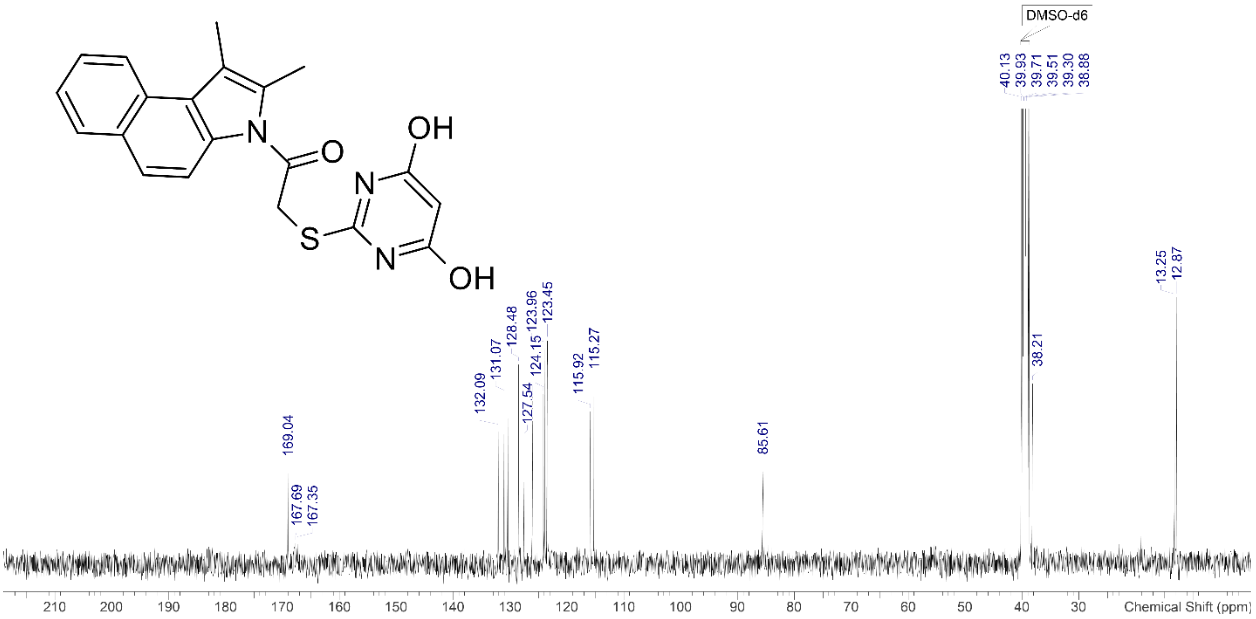


**Figure S20.** $^{1}$H NMR spectrum of compound **12** in $CDCl_3$ (-0.5 – 15 ppm).

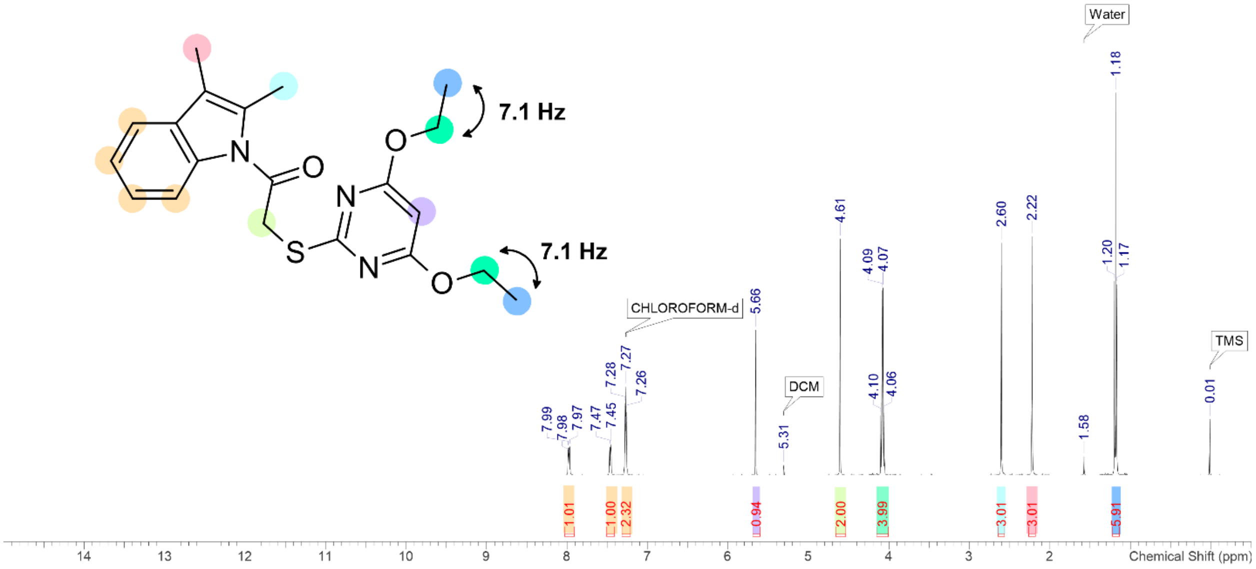

**Figure S21.** $^{13}C$ NMR spectrum of compound **12** in $CDCl_3$ (0 – 220 ppm).

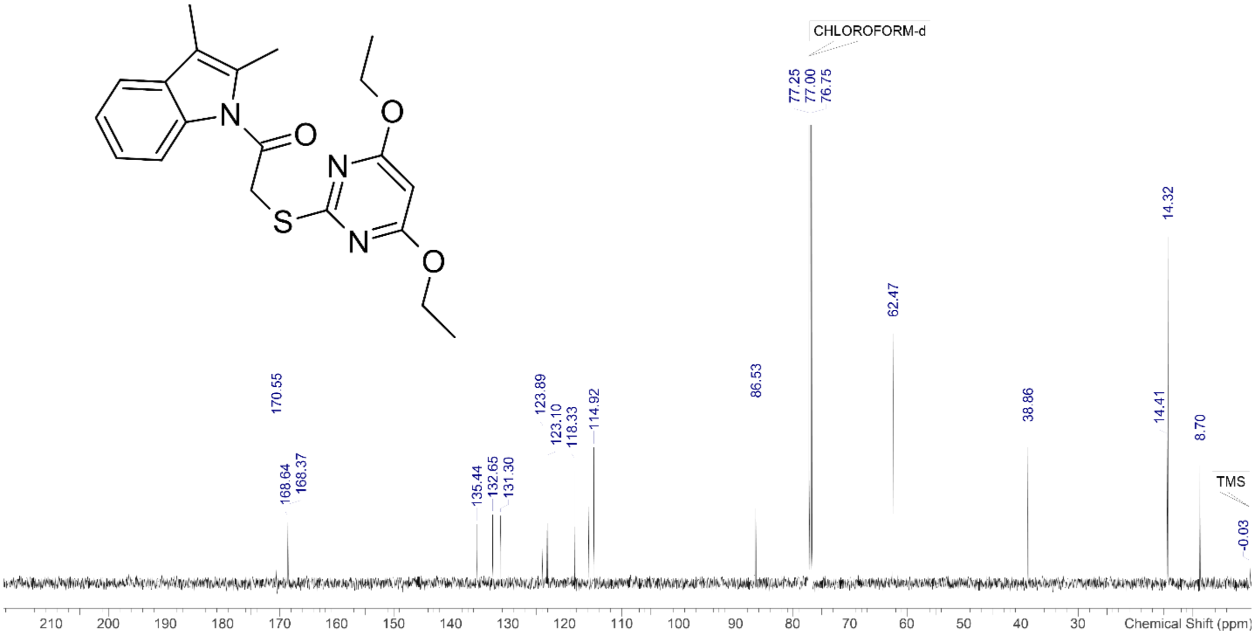


**Figure S22.** $^{1}H$ NMR spectrum of compound **13** in $d_6$-DMSO (-0.5 – 15 ppm).

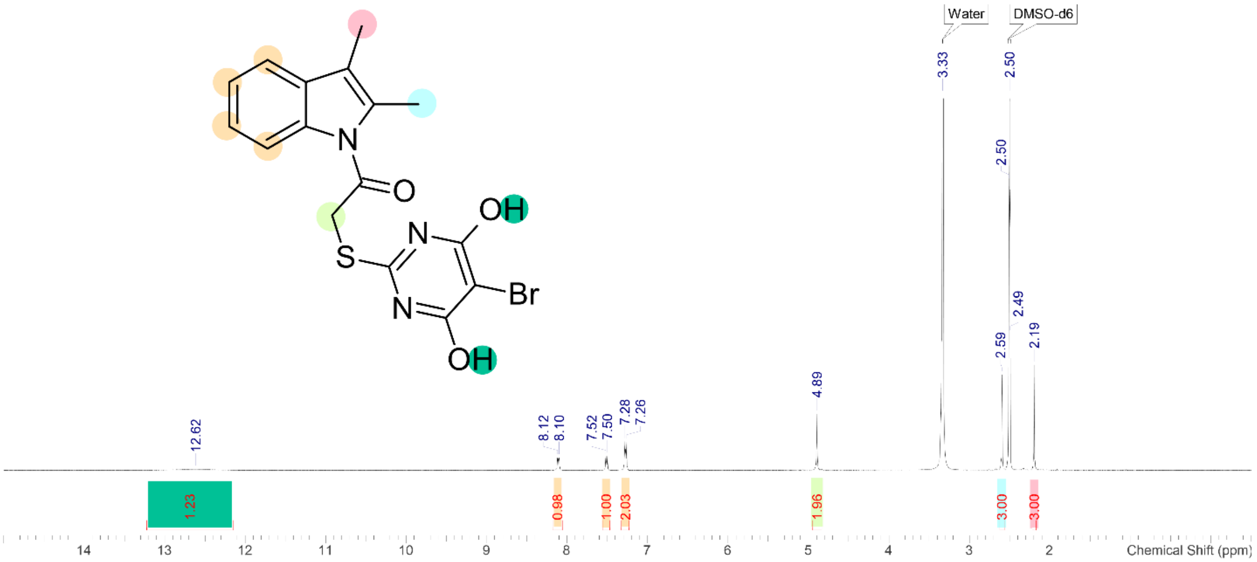

**Figure S23.** $^{13}$C NMR spectrum of compound **13** in $d_6$-DMSO (0 – 220 ppm).

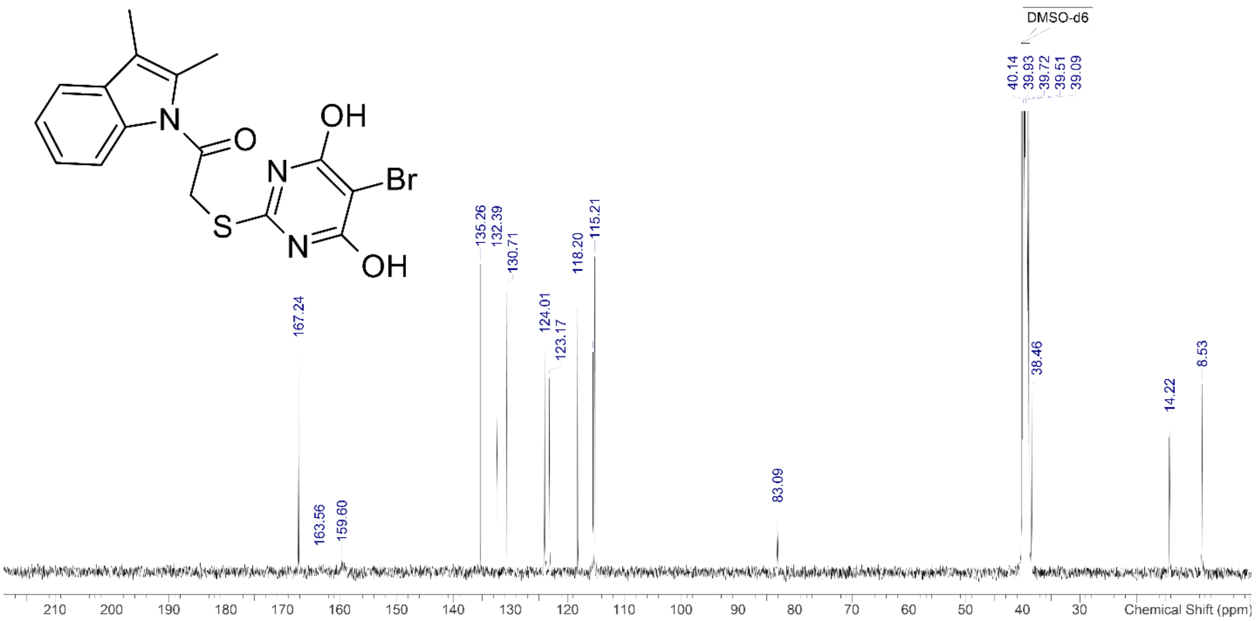
